# UNRAVELing the synergistic effects of psilocybin and environment on brain-wide immediate early gene expression in mice

**DOI:** 10.1101/2023.02.19.528997

**Authors:** Daniel Ryskamp Rijsketic, Austen B. Casey, Daniel A.N. Barbosa, Xue Zhang, Tuuli M. Hietamies, Grecia Ramirez-Ovalle, Matthew Pomrenze, Casey H. Halpern, Leanne M. Williams, Robert C. Malenka, Boris D. Heifets

## Abstract

The effects of context on the subjective experience of serotonergic psychedelics have not been fully examined in human neuroimaging studies, partly due to limitations of the imaging environment. Here, we administered saline or psilocybin to mice in their home cage or an enriched environment, immunofluorescently-labeled brain-wide c-Fos, and imaged cleared tissue with light sheet microscopy to examine the impact of context on psilocybin-elicited neural activity at cellular resolution. Voxel-wise analysis of c-Fos-immunofluorescence revealed differential neural activity, which we validated with c-Fos^+^ cell density measurements. Psilocybin increased c-Fos expression in the neocortex, caudoputamen, central amygdala, and parasubthalamic nucleus and decreased c-Fos in the hypothalamus, cortical amygdala, striatum, and pallidum. Main effects of context and psilocybin-treatment were robust, widespread, and spatially distinct, whereas interactions were surprisingly sparse.

## Introduction

The therapeutic potential of serotonergic psychedelics like psilocybin is promising (Goodwin et al., 2022), but factors contributing to their efficacy are not well understood. It is widely assumed that the environmental setting in which a psychedelic drug is taken shapes the subjective experience (Golden et al., 2022; Hartogsohn, 2017), and some evidence links particular subjective experiences to therapeutic outcomes (Johnson et al., 2019; Roseman et al., 2018). This belief has led clinical researchers to administer psychedelics in controlled environments, often in conjunction with psychotherapy (Davis et al., 2021; Goodwin et al., 2022). However, psychedelic use in uncontrolled, naturalistic environments such as mass gathering events, may similarly elevate mood and promote social connectedness (Forstmann et al., 2020; Nygart et al., 2022). Thus, it is unclear whether psychedelic states, and their underlying neural dynamics, represent two independent effects of drug and setting, or if these factors interact to create setting-specific drug effects. Attempts to understand psychedelic drug action in humans have relied on neuroimaging readouts such as hemodynamic responses and arterial spin labeling derived from functional magnetic resonance imaging (fMRI) (Carhart-Harris et al., 2012; Lewis et al., 2017) and metabolic demand in positron emission tomography (PET) (Gouzoulis-Mayfrank et al., 1999; Vollenweider et al., 1997). However, these modalities are largely limited by confinement of subjects to an imaging environment, leaving unresolved the question of how much setting influences psychedelic brain states.

To study the effect of environmental context on psychedelic-elicited states in mice, we mapped, at cellular resolution, neural activities related to context, psilocybin, and the interaction between these factors. To capture experience-dependent brain activity, we performed immunofluorescent labeling of c-Fos, an immediate early gene (IEG) product and transcription factor that is transiently expressed after neural activation (Sagar et al., 1988), followed by brain clearing via iDISCO+ (Renier et al., 2016) and light sheet fluorescent microscopy (LSFM) (Davoudian et al., 2023; Hansen et al., 2021; Jin et al., 2022; Renier et al., 2016). Psilocybin or saline was administered to mice in two distinct contexts, a familiar home cage or an enriched environment (EE) known to promote neural and behavioral plasticity (Nithianantharajah & Hannan, 2006). While we observed large main effects of context and psilocybin on neural activity across the brain, markedly smaller effects resulted from interactions.

We performed c-Fos mapping in TRAP2 (Targeted Recombination in Active Populations; *Fos*::c-Fos-2A-iCre^ERT2^); Ai14 (*Rosa-CAG-LSL*::tdTomato) mice to determine if identified ensembles are amenable to control by functionally defined genetic recombination (Allen et al., 2017; DeNardo et al., 2019). We identified several brain regions that may be critical to psilocybin’s effects and confirmed that some of these ensembles can be selectively manipulated in future mechanistic studies. Overall, our results favor a model in which acute psychedelic brain states represent the sum of environment and drug effects, with comparatively little contribution from the interaction of drug and environment.

## Results

### UNRAVEL: UN-biased high-Resolution Analysis and Validation of Ensembles using Light sheet images

Mice were administered saline or psilocybin (2 mg/kg, i.p.) as well as 4-OHT (50 mg/kg, i.p.), enabling activity- and Cre-dependent tdTomato expression, just before placement into their home cage or an EE (Figure 1A). After a 2 week washout, mice were administered saline or psilocybin in a counterbalanced fashion, confined to either environment for two hours and then sacrificed to measure c-Fos expression (Chowdhury & Caroni, 2018). Following transcardial perfusion, brains were hemisectioned, immunolabeled, cleared with iDISCO+, and imaged with 3.5 µm isotropic resolution via LSFM (Figure 1B).

**Figure 1:**
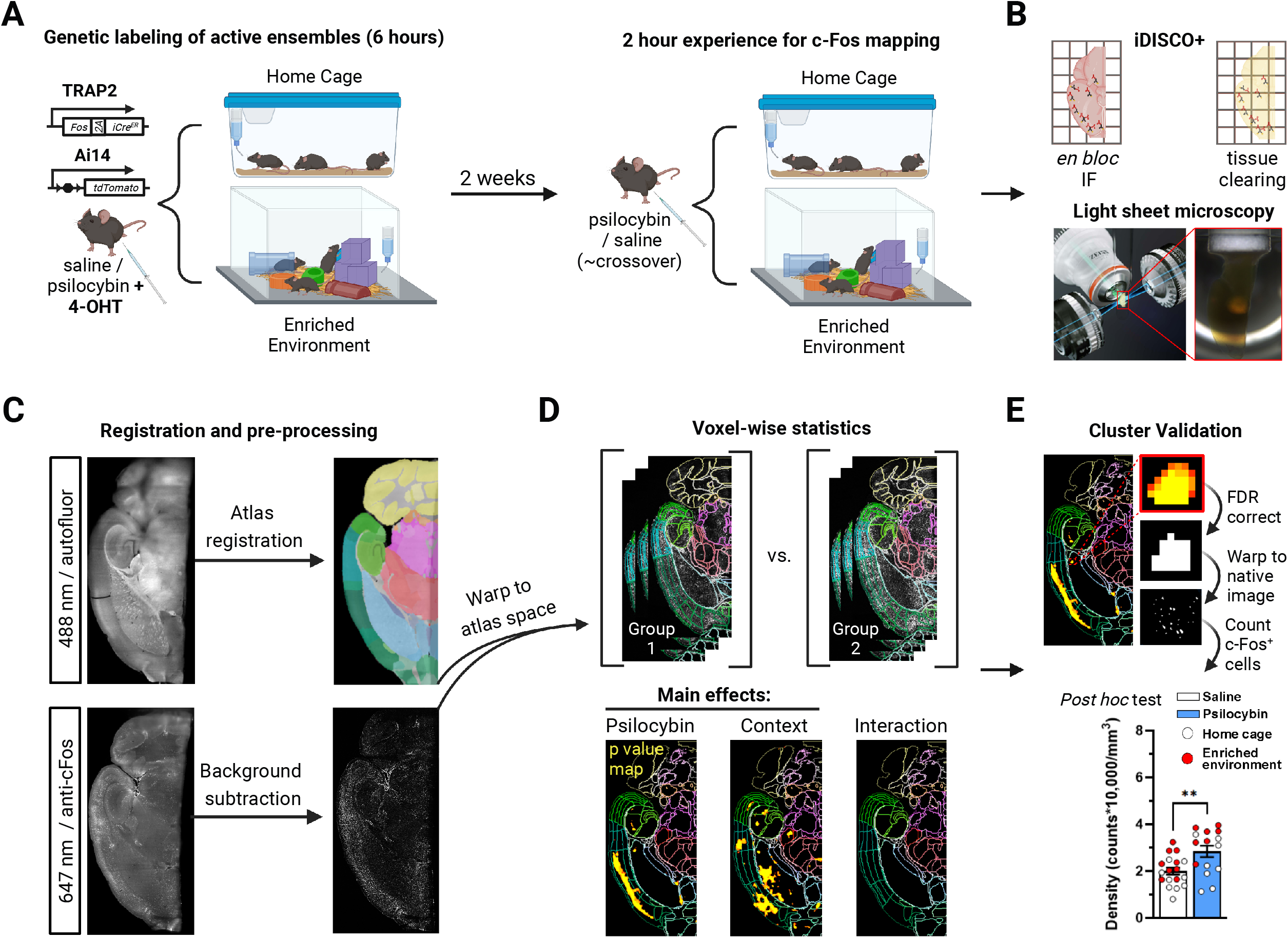
Methodological workflow for the identification and validation of neural populations activated by psilocybin across contexts. A) To genetically label active neurons, mice were injected with psilocybin or saline as well as 4-OHT and placed in their home cage or an enriched environment (EE) with cage mates. To map c-Fos, mice were injected two weeks later with psilocybin or saline and placed in their home cage or an EE with cage mates. B) Brain hemispheres were immunolabeled, cleared, and imaged with LSFM. C) Z-stacks of autofluorescence were registered to a LSFM specific Allen brain atlas. c-Fos immunofluorescence (IF) was rolling ball background subtracted, warped to atlas space using transforms from registration, and z-scored. D) A voxel-wise 2×2 ANOVA identified clusters of voxels with divergent c-Fos-IF intensity. E) Clusters were defined by FDR correction and validated by warping them to tissue space to measure c-Fos^+^ cell densities.

We developed UNRAVEL, an automated command line workflow, to perform voxel-wise analysis of c-Fos immunofluorescence (IF) across the brain and validate identified clusters of significant voxels using c-Fos^+^ cell density quantification at cellular resolution (Figure 1C-E). Registration to an LSFM-based averaged template brain, which is aligned with the Allen brain atlas, was performed using autofluorescence image volumes (Figure 1C). c-Fos-IF image volumes were background subtracted, warped to LSFM atlas space, z-scored, and used as inputs for voxel-wise statistics according to a 2×2 ANOVA design (Figure 1C-D). The three resulting statistical contrasts represented main effects of context and psilocybin treatment, as well as an interaction term.

Filtering out false positives is a formidable issue for human neuroimaging as it often involves ∼100,000 comparisons (Eklund et al., 2016; Woo et al., 2014). Voxel-wise analysis of IF-labeled LSFM-imaged tissue is a relatively new high-resolution technique more prone to false positives due to a greater number of voxel-wise comparisons. Our analysis, performed at 25 µm isotropic resolution, involved ∼8,300,000 comparisons. Many studies involving voxel-wise analysis of LSFM data apply arbitrary cluster-defining thresholds (Woo et al., 2014). We therefore developed a multi-thresholding approach using false discovery rate (FDR) correction to preferentially enhance the detection of true positive clusters. Briefly, across a range of FDR thresholds, we calculated the c-Fos^+^ cell density of each cluster using cellular resolution IF images and performed *post hoc* comparisons to establish a ground truth of cluster “validity” (Figures 1E, 2C, 3C, S1, S2A-F, and S3). We selected the FDR correction threshold (q) that optimized true-positive cluster identification at the highest allowable spatial specificity (q < 0.01 for main effects and q < 0.15 for interactions; Figure S2F). Together, experimental manipulations altered neural activity in valid clusters composed of 209 separate brain regions (Table S1).

### Brain-wide effects of an enriched environment on c-Fos expression

Our voxel-wise analysis of c-Fos-IF indicated a strong main effect of environmental context, represented by 71 discrete clusters of significant voxels (Figures 2A-C and S2A-B; Tables S1-2), with a distribution similar to prior literature (Ali et al., 2009; van Praag et al., 2000). To check if these differences represented a change in c-Fos expression at cellular resolution, *post hoc* unpaired t-tests comparing c-Fos^+^ cell densities between groups for each cluster revealed 40 valid clusters, 37 of which had increased neural activity in the EE context (Figures 2A-C and S2A-B). Neural activity in the EE condition was increased in subregions of the isocortex, hippocampal formation, olfactory cortex, and thalamus (Figures 2A-C and S2A-B; Tables S1-2). Activity was decreased in three clusters localized to the cortical subplate (lateral amygdala and dorsal endopiriform nucleus), caudoputamen, and ventrolateral geniculate complex (Figure 2A-C; Tables S1-2). These results confirm that EE exposure broadly activated cortical and subcortical areas with marginal c-Fos downregulation across the brain.

**Figure 2:**
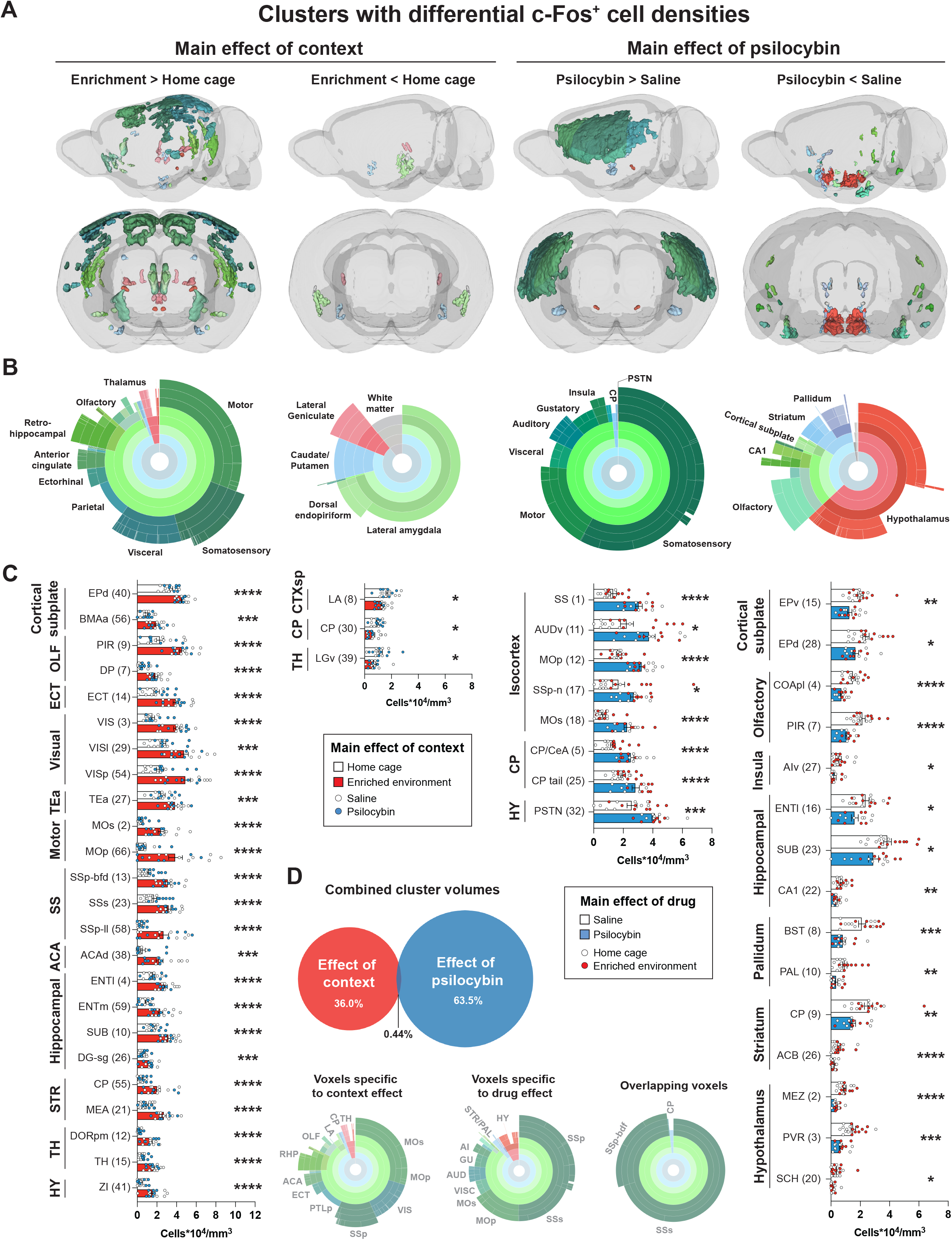
Main effects of context and psilocybin on c-Fos-IF. Clusters with a confirmed difference in c-Fos^+^ cell density are shown. Data is separated in columns based on main effects and effect directions. A) 3D view of clusters from main effects surviving voxel-wise FDR correction (q < 0.01) with Allen brain atlas (ABA) coloring. B) Sunburst plots representing the relative proportion of region volumes for valid clusters at all levels of the ABA (inner rings represent parent regions, whereas outer rings represent the finest level of anatomical granularity). C) *Post hoc* unpaired t-tests of c-Fos^+^ cell density measurements for each cluster. Cluster numbers are in parenthesis. The largest component region of each cluster is indicated. Additional significant and non-significant clusters are presented in Figure S2A-D. D) Relative volumes of main effects, overlap, and corresponding regional volumes. Abbreviations are defined in Table S1. Raw densities and p values for all clusters are in Table S2. \**p* < 0.05, ** p < 0.01, ***p < 0.001, ****p < 0.001.

### Brain-wide effects of psilocybin on c-Fos expression

A robust main effect of psilocybin treatment on brain-wide c-Fos expression was represented by 33 clusters spanning cortical and subcortical structures. *Post hoc* comparisons yielded 27 valid clusters with bidirectional effects on c-Fos^+^ cell density, independent of context (Figures 2A-C and S2C-D; Tables S1-S2). Among the validated clusters, 8 showed a significant increase in c-Fos^+^ cell density (Figure 2A-C). Regional volume measurements indicated that psilocybin-stimulated activity primarily localized to neocortical structures including somatosensory, motor, visceral, auditory, gustatory, and insular cortices. Sparse increased subcortical activity was observed in distinct areas of the caudoputamen, central amygdala, and parasubthalamic nucleus (Figures 2A-C and S2C; Tables S1-2). Decreased activity was found in 19 clusters, present mostly in subcortical areas including the hypothalamus, striatum, and pallidum as well as olfaction-related cortical plate areas including cortical amygdalar areas and piriform areas. Activity was also decreased in CA1 and the endopiriform nucleus (Figures 2A-C and S2C-D; Tables S1-2). Together, our data suggest that psilocybin primarily enhances neural activity in the neocortex while suppressing activity in subcortical regions.

Among voxels surviving FDR-correction and validation, only 0.44% were shared between context- and drug-effects. Overlapping voxels corresponded to secondary somatosensory cortex (context clusters 13 and 63; psilocybin cluster 1), the barrel field of the primary somatosensory cortex (context clusters 22, 25, 33, and 35; psilocybin cluster 1), and the tail of the caudoputamen (context cluster 63; psilocybin cluster 25; Figure 2D).

For valid clusters in which psilocybin increased activity, we tested if the TRAP2 system could be used to label these ensembles, for future behavioral manipulations requiring genetic access. Therefore, using the contralateral hemisphere from the brains of TRAP2^+/-^;Ai14^+^ mice used in c-Fos mapping, we quantified tdTomato^+^ cell densities within the 8 psilocybin-stimulated activity clusters identified above. We found that psilocybin treatment significantly increased tdTomato^+^ cell densities in neocortical cluster 1, striatoamygdalar cluster 5, as well as clusters 11 (layers 4/5 of auditory and temporal association areas) and 18 (secondary motor cortical layers 2/3) (Figure S3). These findings suggest that TRAP2 mice can be used for genetic access to at least a portion of psilocybin-activated neural ensembles.

### Interactions between psilocybin and environmental context

A primary goal of our study was to test for the presence of a context × treatment interaction. That is, we designed our ANOVA comparison to determine if the effect of one variable (psilocybin) on neural activity depends on another variable (the context under which psilocybin was administered). Therefore, we generated a voxel-wise statistical contrast of the interaction term. Seven clusters survived FDR-correction (q < 0.15; Table S2) and *post hoc* 2×2 ANOVA analyses on c-Fos^+^ cell density indicated the presence of 2 valid interactions—in the ventral anterior cingulate cortex and dorsal taenia tecta (Figure 3A-C; Tables S1). In these clusters, psilocybin blocked the c-Fos stimulating effect of the EE (Figure 3C). Voxels comprising valid interaction clusters shared 19% overlap with valid voxels associated with main effects. This overlap occurred in the dorsal taenia tecta (context cluster 7; interaction cluster 3). Conversely, 0.05% of valid main effect voxels overlapped with valid interaction voxels. Consistent with a paucity of significant interaction clusters (Figure 2A vs. Figure 3A), main effect maps had more voxels with larger effect sizes than the interaction map (Figure 3D).

**Figure 3:**
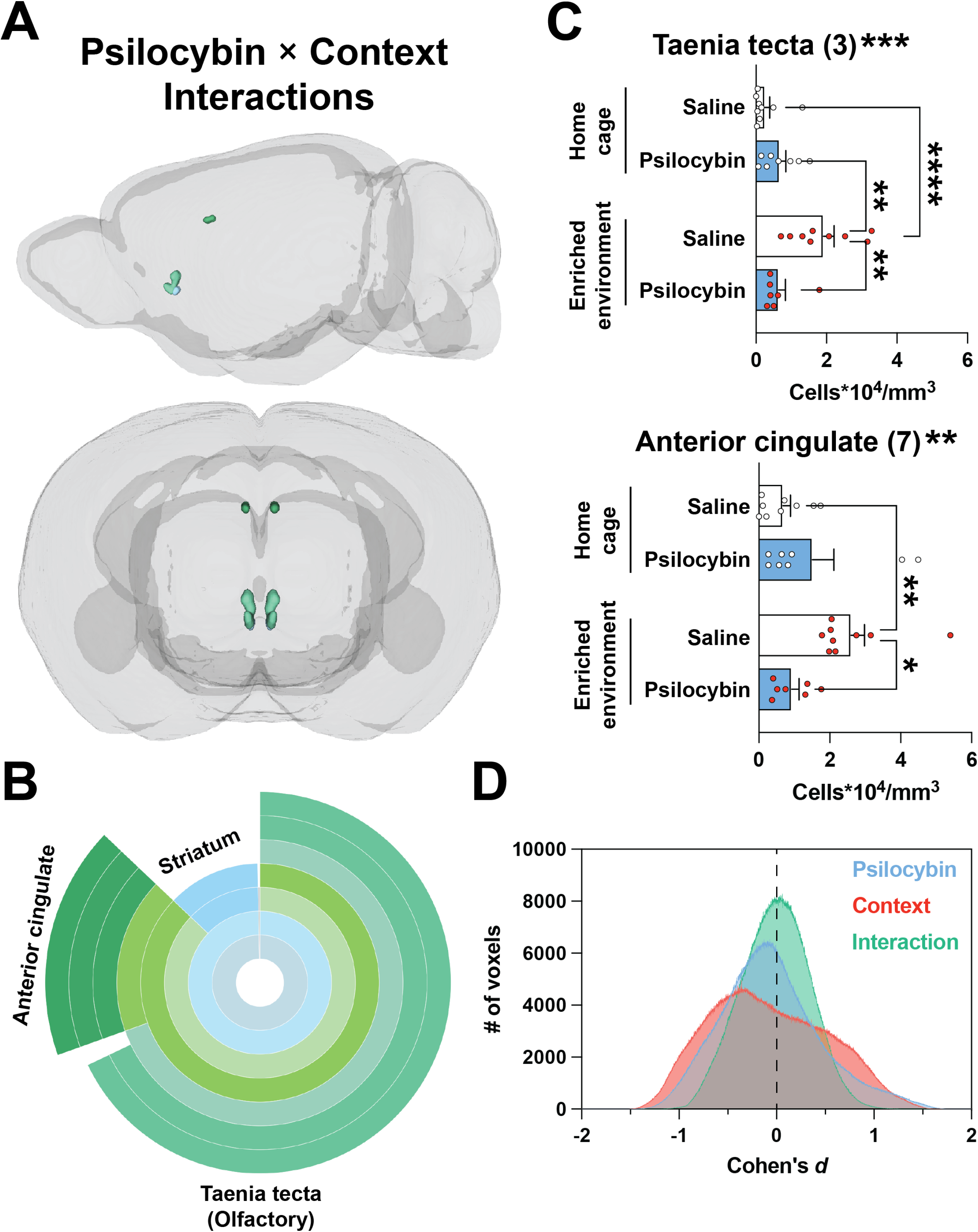
Modest interactions between context and psilocybin. Clusters are shown that survived voxel-wise FDR correction (q < 0.15) and had a confirmed interaction based on *post hoc* 2×2 ANOVAs of c-Fos^+^ cell density measurements. A) 3D view of valid interaction clusters. B) Sunburst plot representing regional volumes. C) *Post hoc* 2×2 ANOVAs of c-Fos^+^ cell densities. Interaction significance is above the graphs. See Figure S2E for non-significant clusters. D) The effect size for each voxel for each contrast is plotted as a histogram binned in steps of Cohen’s *d* = 0.001. \**p* < 0.05, ** p < 0.01, ***p < 0.001, ****p < 0.001.

To extend our search for a context × treatment interaction, we reanalyzed the c-Fos^+^ cell density data shown in Figure 2 using a *post hoc* 2×2 ANOVA for each cluster. Our reanalysis indicated that a significant interaction was present in 5 out of 27 valid psilocybin clusters: striatoamygdalar cluster 5, pallidal cluster 10, auditory cluster 11, dorsal endopiriform nucleus cluster 28, and parasubthalamic nucleus cluster 32. Moreover, a significant interaction was also present in 5 out of 40 valid context clusters: motor cluster 2, dorsal peduncular cluster 7, somatosensory cluster 23, dorsal endopiriform nucleus cluster 31, and dorsal anterior cingulate cluster 38. In general, psilocybin attenuated, or did not change, the c-Fos stimulating effect of the EE. In cases where psilocybin increased activity, the effect was more robust in the familiar home cage condition, whereas psilocybin more effectively decreased activity in the novel EE.

### Psilocybin alters co-activity patterns and modularity of valid clusters

To better understand the impact of psilocybin on brain activity patterns, we performed network analyses on the 27 valid clusters sensitive to psilocybin across environmental contexts. Correlation matrices of c-Fos^+^ cell density yielded a distinct pattern of co-activity between cluster pairs in saline- and psilocybin-treated groups (Figure 4A-B). Notably, although 19 clusters were associated with decreased activity after psilocybin administration, no anticorrelated cluster activity was significant. To visualize differences between cluster co-activity across treatment conditions, we subtracted Pearson correlations in the saline group from those in the psilocybin group (Figure 4C), such that correlated activity common across treatment groups have values near 0. The top 5% of differences in co-activity between groups were primarily represented by cluster pairs with highly correlated activity in the saline group that dissipated following psilocybin treatment.

**Figure 4:**
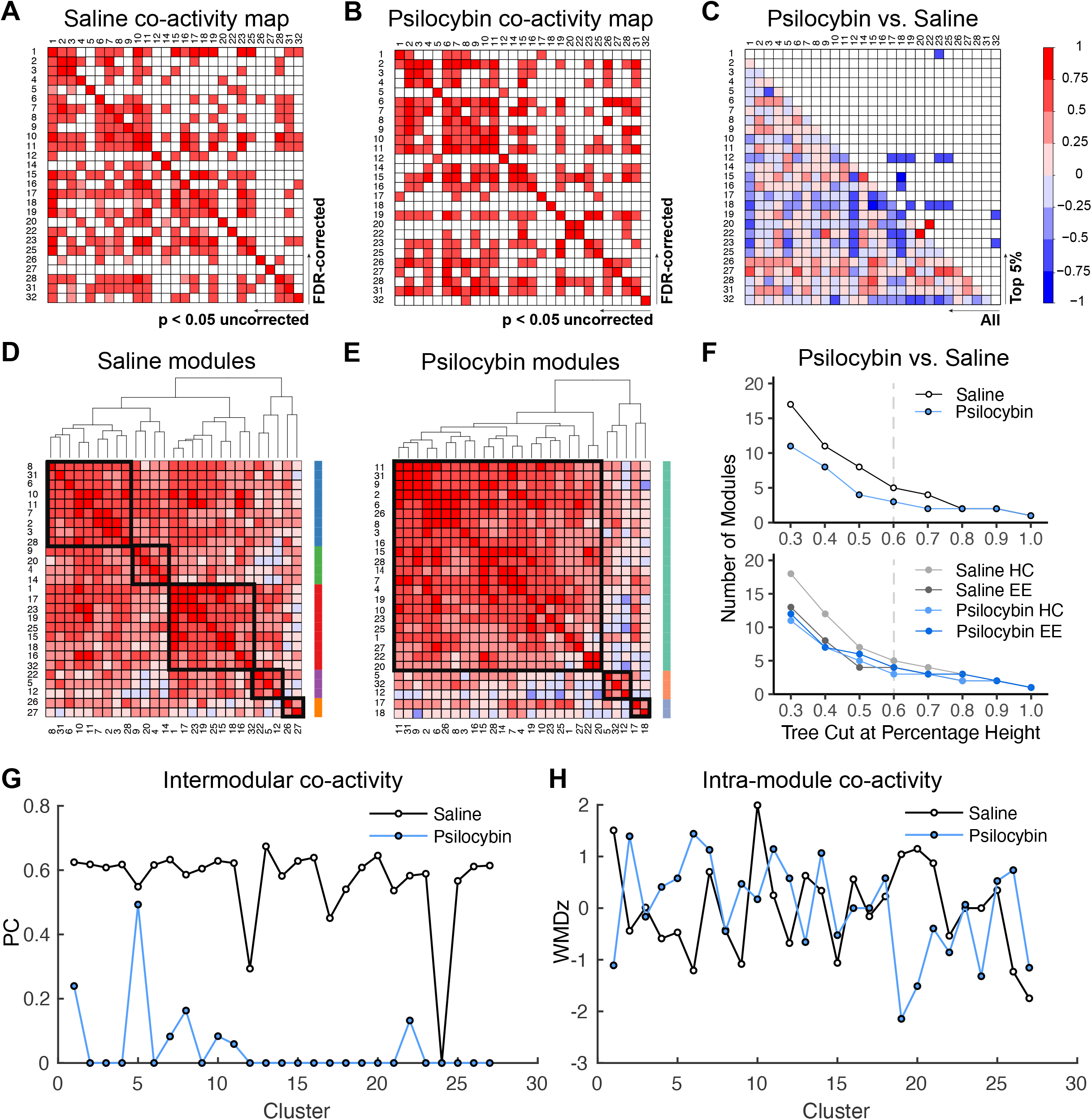
Network analysis of neural populations sensitive to psilocybin. A,B) Pearson correlation matrix of c-Fos^+^ cell densities, in valid clusters where psilocybin altered activity, following treatment of A) saline or B) psilocybin. FDR-corrected (q < 0.05) and uncorrected correlations are above and below the diagonal, respectively. C) Difference map of the Pearson R coefficients. The color scale represents the degree of correlation (Pearson’s r) for each cluster pair. D,E) Hierarchical clustering of uncorrected correlation coefficients at a tree height of 60% indicated the presence of D) five distinct modules in saline-treated brains and E) three modules in psilocybin-treated brains. Tree-based representations of hierarchical clustering are outside of the correlograms. Black squares in the correlograms and the colored bars demarcate co-activity modules. F) Dendrogram cutting at different tree heights indicates the stability of decreased modularity in the psilocybin-treated cohort. The dashed line represents the selected tree height. G) Intra- and H) inter-modular co-activity for each cluster for saline and psilocybin conditions.

We then systematically defined group differences in correlated activity by exploring the modular organization of psilocybin-sensitive neural populations via hierarchical clustering of Pearson correlations based on complete Euclidean distances. The resulting modules reflect cluster networks with highly correlated activity. When trimming at 60% of tree height, saline organized the pattern of co-activity into 5 distinct modules (Figure 4D), whereas psilocybin restructured the pattern of co-activity to give only 3 modules (Figure 4E). The psilocybin-mediated decrease in network modularity was independent of hierarchical cluster-thresholding, and this effect was driven by greater modularity in mice treated with saline in the home cage (Figure 4F). Brains from mice treated with saline in the EE exhibited a similar number of modules to those treated with psilocybin, independent of context. Although within-module co-activity was similar for saline and psilocybin-treated groups (Figure 4G), substantially less between-module co-activity was observed in psilocybin-treated mice (Figure 4H).

## Discussion

Psilocybin therapy is a promising rapid-acting intervention for relief from some types of major depressive disorder and similar negative affective states in substance use disorders. The subjective effects of psilocybin and its putative therapeutic efficacy may depend on the environmental context of drug administration (Carhart-Harris et al., 2018). There are remarkably few studies, in any species, that systematically study how this ostensibly crucial non-drug parameter impacts the subjective effects, neurophysiology, or therapeutic outcomes associated with serotonergic psychedelics like psilocybin (Carhart-Harris, 2023). Human neuroimaging techniques are not amenable to major environmental manipulations (e.g., those involving naturalistic movement) and are therefore limited at exploring contextually specific, psilocybin-elicited neural activity. We posed this question in a more tractable mouse model using a broad environmental manipulation (home cage versus enriched environment) that does not assume which mouse behavioral outcomes are therapeutically relevant, and with a psilocybin dose (2 mg/kg, i.p.) that falls within the range widely used across mouse studies (Davoudian et al., 2023; Fadahunsi et al., 2022; Golden & Chadderton, 2022; Hesselgrave et al., 2021; Shao et al., 2021).

Advances in preclinical neuroimaging allowed us to directly test the role of environmental setting on psilocybin-elicited neural activity across the mouse brain using an unbiased analysis pipeline combining LSFM, human neuroimaging toolkits, and custom scripts (Figure 1; Figure S1). Mice were injected with saline or psilocybin in a familiar home cage context or a novel EE, and experience-dependent neural activity was quantified at cellular resolution throughout the brain. Voxel-wise analyses revealed strong main effects of EE or psilocybin exposure on neural activity defined by spatially distinct clusters of differential c-Fos-IF. Since voxel-wise statistics are prone to false positives (Eklund et al., 2016), we implemented FDR-correction and validated clusters in full resolution images using *post hoc* tests of c-Fos^+^ cell density. In sum, 40 and 27 valid clusters represented neural populations sensitive to EE and psilocybin treatment, respectively. Two small clusters corresponded to valid context × psilocybin treatment interactions. Contrary to popular belief, these findings suggest that the psilocybin experience is primarily driven by the large and independent effects of environment and the drug, rather than a synergistic effect of these factors.

Our novel analytical approach, UNRAVEL (Figure 1), allows for robust detection and validation of spatially restricted changes in c-Fos-IF or the intensity of other labels. UNRAVEL incorporates several innovations in the analysis of LSFM data from our group and others.

MIRACL accurately registers cleared brain tissue (Goubran et al., 2019), but uses an average template brain from serial two-photon microscopy that has an autofluorescence intensity profile distinct from our samples. Thus, we improved registration accuracy by supplementing MIRACL with an averaged template brain generated using iDISCO+ and LSFM (Perens, Salinas, et al., 2021). Importantly, we implemented a voxel-wise analysis method instead of regional analysis, allowing us to detect focal differences in signal either occurring within a brain subregion not specified by standard atlases or crossing anatomical demarcations. This enhanced anatomical resolution comes at the cost of false positive discovery. Studies implementing voxel-wise analyses often apply arbitrary thresholds for multiple comparisons correction, if a correction is applied at all (Woo et al., 2014). UNRAVEL comprehensively characterizes the accuracy of corrected activity maps based on quantification of cell densities within clusters surviving correction across a range of FDR thresholds, allowing for the selection of an optimal q value. As part of cluster validation, cell segmentation was democratized by a consensus approach wherein voxels were preserved if mutually classified as c-Fos^+^ by a majority of raters. Moreover, we simplified qualitative validation by automating extraction of raw, background subtracted, and segmented images corresponding to the most significantly different portion of each cluster (Figure S1). Our approach enabled a high resolution, brain-wide analysis of two powerful factors influencing brain activity.

Environmental enrichment has been studied intensely due to its therapeutic-like influence on neural plasticity and behaviors associated with autism spectrum disorder and substance abuse in animal models (Baroncelli et al., 2010; Ey et al., 2011; Solinas et al., 2008; van Praag et al., 2000). We observed that acute exposure to an EE broadly enhanced c-Fos-IF across cortical and subcortical areas, despite pooling saline and psilocybin conditions to measure the main effect of context. Consistent with reports that an EE increases voluntary exercise in rodents (van Praag et al., 2000), we observed increased c-Fos^+^ cell density in motor cortices. The EE also increased c-Fos-IF in the dentate gyrus (granule cell layer) and lateral amygdala (context clusters 26 and 8, respectively), both of which reportedly increase activity after a single EE exposure (Ali et al., 2009). Moreover, cFos-IF was increased in layer 5 of the lateral entorhinal cortex (context cluster 4), which may contribute to novel object discrimination in rodents (Wilson et al., 2013). Together, our results suggest that EE increased c-Fos^+^ cell density in regions associated with motor behavior, context recognition, and novelty.

In contrast, independent of context, psilocybin diffusely upregulated c-Fos expression across the neocortex, including somatosensory, motor, visceral, auditory, gustatory, agranular insular, and temporal association cortices. The bulk of increased c-Fos-IF corresponded to a single continuous cluster (psilocybin-treatment cluster 1) spanning the anterior-posterior axis along layers 2/3 (∼8%), 4 (∼24%), 5 (∼60%), and 6a (∼8%) of the aforementioned cortices. Increased activity under psilocybin was also observed in comparatively smaller clusters located in discreet neocortical areas (clusters 11, 12, 17, and 18), striatoamygdalar areas (cluster 5), the tail of the dorsal striatum—which may facilitate hallucination-like perceptions (Schmack et al., 2021)—(cluster 25), and the parasubthalamic nucleus (cluster 32). Suppression of neural activity was primarily localized to subregions of the hypothalamus, olfactory areas in the cortical plate, hippocampal formation, cortical subplate, striatum, and pallidum.

Our findings generally agree with a recent regional analysis of psilocybin-mediated c-Fos expression across the mouse brain using LSFM and two-photon microscopy (Davoudian et al., 2023). However, our higher resolution voxel-wise analysis technique resolved focal changes in neural activity among more sizeable structures that may have modest or no changes in activity when considered as a whole. Moreover, the resolution and specificity obtained with UNRAVEL allowed us to detect heterogeneous activity within specific brain regions. For example, we detected a small neural ensemble in the parasubthalamic nucleus that was activated by psilocybin (cluster 32), despite this region having decreased activity when considered as a whole (Davoudian et al., 2023), thus emphasizing the utility of our method to guide local stereotactic injections with the goal of further understanding experience-dependent neural activity.

Our results on psilocybin-elicited neural activity in mice parallel findings in humans measured by glucose metabolism (PET) or blood flow (fMRI). In healthy human volunteers, psilocybin diffusely increased glucose metabolism in frontal cortical areas and decreased metabolism in subcortical areas such as the pallidum (Gouzoulis-Mayfrank et al., 1999; Vollenweider, 1998; Vollenweider et al., 1997). FMRI studies noted decreased blood flow in the hypothalamus, striatum, putamen, and hippocampus after psilocybin administration (Carhart-Harris et al., 2012; Lewis et al., 2017). However, such human brain imaging studies have not been able to define the influence of “setting” (i.e., the environment) on psilocybin-induced brain-wide neural activity. Surprisingly, we found that psilocybin-mediated changes in neural activity are largely context-independent. In detected clusters representing a drug × setting interaction, psilocybin suppressed EE-driven c-Fos enhancement. While sparse, these regions may be suitable for probing context-specific behavioral effects. For example, interaction cluster 7 localized to the ventral part of the anterior cingulate cortex, a region associated with context-dependent cue-reinstatement of cocaine responding (Torregrossa et al., 2013) and fear memory generalization (Cullen et al., 2015) in rodents as well as threat response regulation in humans (Williams, 2017).

Our dataset also allows a cross-species comparison of functional connectivity under the influence of psychedelics. Measuring correlations of variance in c-Fos^+^ cell densities between seed clusters and hotspots revealed that psilocybin changed the distribution of co-active clusters while decreasing modularity and intermodular co-activity. These changes in network activity are reminiscent of the disintegration of functional connectivity within association networks caused by psilocybin in several human studies (Kwan et al., 2022). This decoupling may reflect a hallmark subjective effect of psilocybin, perhaps produced by affecting the integration of interoceptive and exteroceptive stimuli in a context-independent manner. This apparently conserved physiology could form the basis for future work testing the causal influence of these network properties on behavioral changes associated with psychedelic use.

The use of c-Fos as a proxy of neural activity has limitations in that it provides low temporal resolution (∼5 hours) (Chowdhury & Caroni, 2018; Morgan et al., 1987) and not all cells are equally c-Fos competent (Hoffman et al., 1993; Sgambato et al., 1997). Additionally, some mice in our study received psilocybin two weeks prior to c-Fos mapping (Figure 1A,B), which may limit the interpretability of our data if psilocybin persistently modified neural dynamics over that time frame, a possibility which is difficult to evaluate as most rodent studies do not report changes in dendritic structure, electrophysiological properties, or behavior past 7 days (Hesselgrave et al., 2021; Hibicke et al., 2020; Shao et al., 2021). Despite the biological variation contributed by animal sex, genotype, and data pooling, we observed robust main effects of psilocybin-elicited c-Fos expression (Voelkl et al., 2020).

While human neuroimaging studies provide valuable insights, they neither capture the full impact of context on drug-induced brain activity, nor do they establish clear causal relationships between specific neurophysiological events and therapeutic outcomes. Our results in mice suggest that both environmental context and psilocybin each elicit independent and pronounced brain-wide neural responses, which are primarily additive rather than synergistic. Furthermore, our demonstration that the TRAP2 mouse line can be used to genetically capture, and thereby presumably control, neural ensembles activated by psilocybin, provides a platform for determining the networks and cell types that mediate psilocybin’s therapeutic-like effects and whether these effects are context- or state-dependent.

## Supporting information

3D neuroimaging data

## Acknowledgments

The authors offer their sincere gratitude to Dr. Michel B. Hell for his help with adapting the CLIJx plugin to allow for fractional assignment of cells overlapping region boundaries during 3D object counting on the GPU, Mr. Daniel F. Cardoza Pinto for helpful discussions, and Miss Zahra Rastegarmoghaddam for her assistance with initial adaptation of an excel template for use with the LSFM atlas. We also thank the Wu Tsai Neuroscience Center, Neuroscience Microscopy Service for initial guidance on light sheet microscopy and help collecting pilot data. Figure 1 was created with BioRender. XZ and LMW acknowledge support from the National Institute of Drug Abuse under award P50DA042012.

## Author Contributions

**D.R.R**.: Writing – original draft, Conceptualization, Investigation, Data Curation, Formal analysis, Methodology, Software, Validation, Visualization.

**A.B.C**.: Writing – original draft, Conceptualization, Investigation, Data Curation, Formal analysis, Methodology, Software, Validation, Visualization.

**D.A.N.B**.: Conceptualization, Methodology, Software, Writing - Review & Editing.

**X.Z**.: Formal analysis, Methodology, Software, Visualization, Writing - Review & Editing.

**T.M.H**.: Conceptualization, Writing - Review & Editing.

**G.R.O**.: Software.

**M.P**.: Conceptualization, Methodology, Writing - Review & Editing.

**C.H.H**.: Writing - Review & Editing, Supervision.

**L.M.W**.: Writing - Review & Editing, Supervision.

**R.C.M**.: Writing - Review & Editing, Supervision.

**B.D.H**.: Writing - Review & Editing, Conceptualization, Methodology, Funding acquisition, Project administration, Resources, Supervision.

## Methods

### Animals

Experiments were designed to address two goals: mapping the brain-wide distribution of c-Fos and, in the same mice, establishing the feasibility of controlling neural ensembles activated by psilocybin using the TRAP2 mouse line. In TRAP2 mice (*Fos*::c-Fos-2A-iCre^ERT2^ knock-in; RRID:IMSR_JAX:030323), c-Fos and iCre^ERT2^ are expressed in cells with *Fos* promoter activity and separated via ribosomal skipping of the 2A coding sequence during translation. Injection of 4-hydroxytamoxifen (4-OHT) activates iCre^ERT2^, enabling translocation from the cytosol to the nucleus for site-specific recombination. TRAP2 mice were crossed to an Ai14 reporter line (*Rosa-CAG-LSL*::tdTomato; RRID:IMSR_JAX:007914) (Allen et al., 2017) for visualization of activated neurons via conditional tdTomato expression. Prior to the TRAPing timepoint, mice were naïve to experiments and treatments.

Twenty-five male and 8 female TRAP2;Ai14 mice (12–18 weeks of age) were bred in house to C57BL6/J wild type mice to give littermates of TRAP2^+/-^ or TRAP^-/-^, and Ai14^+/+^, Ai14^+/-^ or Ai14^-/-^ genotypes (Transnetyx; Cordova, TN); a subset of mice (16 males and 6 females, TRAP^+/-^;Ai14^+^) was used to assess the feasibility of genetically labeling psilocybin-activated neural populations. Animals were maintained in age- and sex-matched groups of 2–5 littermates and housed in an 12:12 light-dark cycled SPF facility and appeared to be in good health; male mice weighed 31.4 ± 0.7 g (mean ± SEM) and females weighed 23 ± 0.9 g. Home cages were sterile ventilated cages by Innovive (San Diego, CA) offering irradiated 1/8” corn cob bedding (The Andersons, Inc.) with *ad libitum* access to pre-filled acidified water bottles (Innovive) and irradiated 18% protein rodent diet (Teklad Global). All animal behavioral procedures comply with the ARRIVE guidelines (Sert et al., 2020), the Guide for the Care and Use of Laboratory Animals (Council, 2011), and were approved by the Stanford University Institutional Animal Care and Use Committee. The animal care and use program is fully accredited by AAALAC, International and holds an Assurance with OLAW.

### Habituation and drug administration

Mice were habituated to handling for 5 minutes in the 3 days prior to experimental manipulations. On test days, mice in the home cage context remained in the vivarium, whereas mice exposed to an EE were transported to a dimly lit neighboring experimental room kept at 22 °C. Mice were habituated for at least 60 minutes before handling. During habituation, psilocybin was diluted to 0.2 mg/mL in sterile saline from 100 mM stocks in DMSO (final [DMSO] ≤ 1%), and 4-OHT was prepared in corn oil at 5 mg/mL. All injections were performed i.p. at 0.1 mL/10 g body weight. Animal handling and injections were performed by a male experimenter (D.R.R). Psilocybin was acquired from the Research Triangle Institute (Durham, NC) through the National Institutes on Drug Abuse drug supply program.

### Stimulation and capturing of mouse neural activity

Mice were divided into four groups and injected with saline or a behaviorally active dose of psilocybin (2 mg/kg) (Shao et al., 2021) in a familiar home cage context or a novel EE. The home cage context consisted of standard vivarium housing described above, whereas the EE was assembled using a larger transparent cage (Innovive, product #: R-BTM-H), sawdust bedding, a tube, and 5 novel objects which were varied to retain novelty. Food and water were available *ad libitum*, and treatment conditions were consistent across cage mates.

The first exposure to saline or psilocybin was followed immediately by an injection of 4-OHT (50 mg/kg) to label activated cells in TRAP2^+/-^;Ai14^+^ mice. Mice treated in their home cage were left on the vivarium rack, whereas mice treated in the EE were returned to their home cage 6 hours later. All mice were then left undisturbed in the vivarium for a two-week washout period, save for standard animal husbandry such as general health inspections and cage changes. After the washout period, mice were injected again with saline or psilocybin (in the absence of 4-OHT), according to a crossover design, and allowed to roam their assigned cage for 2 hours before sacrifice.

### Perfusions and sample preparation

Mice were sacrificed by perfusion under isoflurane anesthesia. Perfusions were initiated with 10 mL of 1x PBS (pH=7.4, rt), followed by 10 mL of 4% paraformaldehyde in 1x PBS (pH=7.4, rt) at a rate of ∼3 mL/min. Brains were extracted and incubated overnight at 4 °C in 10 mL of 4% paraformaldehyde in 1x PBS. The next day, brains were brought to room temperature (rt), washed 3x 30 min with 10 mL of 1x PBS supplemented with 0.02% (w/v) sodium azide (pH = 7.4, rt) while gently rocking on a nutator (Fisherbrand), olfactory bulbs were removed, and whole brains were manually cut into hemispheres with a razor blade. Samples were stored at 4 °C in 10 mL of 1x PBS + 0.02% sodium azide (pH = 7.4, 4 °C) until continuing with the IF labeling/iDISCO+ procedure described below.

### Immunofluoresent (IF) staining and clearing

The iDISCO+ procedure was performed as described (Renier et al., 2014) with slight modifications. Brain hemispheres were brought to rt and washed 3x 30 min with 2 mL of 1x PBS + 0.02% sodium azide (pH = 7.4, rt) while revolving at low speed on a tube revolver (ThermoScientific). Samples were dehydrated, bleached, and rehydrated, to remove membrane lipids, enhance permeabilization and reduce autofluorescence (Renier et al., 2014). For this, each sample then underwent five sequential 45 min rounds of dehydration in 2 mL of: 20%, 40%, 60%, 80%, and 100% methanol in 1x PBS (v/v, pH=7.4, rt) while revolving. A final 1 hr dehydration step in 100% methanol was performed before chilling samples to 4 °C and incubated overnight at rt in 2:1 dichloromethane:methanol under constant revolution. The next day, samples were washed 2x 1 hr with 100% methanol (rt) while revolving, chilled to 4 °C, and incubated overnight at 4 °C in 2 mL of 5% hydrogen peroxide (1:5 water:methanol). After bringing samples to rt, samples were rehydrated with five sequential 45 min incubations in 2 mL of: 80%, 60%, 40%, 20%, and 0% methanol in 1x PBS (v/v, pH=7.4, rt) with constant revolution.

For immunostaining, membranes were permeabilized with 2 mL of 1x PBS + 0.2% Triton X-100 (v/v) while revolving for 1 hr (rt), followed by a 48 hr incubation at 37 °C while revolving in 2 mL of permeabilization solution (0.2% triton X-100 [v/v], 0.02% sodium azide [w/v], 20% DMSO [v/v], and 306 mM glycine). After decanting the permeabilization solution, each sample was incubated 2x 5 min at rt in 2 mL of PTwH (0.2% tween-20 [v/v], 0.01 mg/mL heparin, and 0.02% sodium azide [w/v] in 1x PBS, pH=7.4, rt) while revolving. Samples were blocked by incubating each sample in 1x PBS (pH=7.4) with 0.2% triton X-100 (v/v), 6% donkey serum (v/v), and 10% DMSO (v/v) for 48 hours at 37 °C. Primary antibodies were diluted 1:500 (rabbit polyclonal anti-c-Fos; Synaptic Systems, item #: 226003) or 1:250 (rabbit anti-RFP; Rockland, item #: 600-401-379) in PTwH supplemented with 5% DMSO (v/v) and 3% donkey serum (v/v). Anti-c-Fos and anti-RFP immunolabeling was performed on the left and right hemisphere of a given brain sample (with few exceptions), and immunolabeling was carried out in 1.6 ml for 10 days at 37 °C under constant rotation. After washing 5x 5 min and overnight at rt in 2 mL of PTwH, samples were protected from light going forward and incubated for 5 days at 37 °C in 1.6 ml of PTwH + 3% donkey serum containing a 1:250 dilution of secondary antibody (donkey anti-Rabbit Alexa 647, ThermoFisher Scientific). Samples were washed 5x 5 min and incubated overnight at rt in 2 mL of PTwH.

Tissue clearing was achieved by dehydration with five sequential 45 min incubations in 2 mL of: 20%, 40%, 60%, 80%, and 100% methanol (v/v in 1x PBS, pH=7.4), followed by a 1 hr incubation in 100% methanol, and overnight incubation in 2:1 dichloromethane:methanol (all rotating at rt in the dark). After a final series of washing with 100% dichloromethane (2x 2mL, rotating at rt for 15 min) to remove excess methanol, samples were cleared while sitting still overnight in 2 mL of dibenzyl ether. In total, there were 33 anti-c-Fos labeled hemispheres and 22 anti-RFP labeled hemispheres (n=7–9 mouse hemispheres per group for anti-c-Fos and n=4–8 per group for anti-RFP). Unequal group sizes resulted from sample exclusion prior to analysis due to genotype, or poor tissue or registration quality, though sample sizes used for activity mapping are comparable to other published work (DeNardo et al., 2019; Hansen et al., 2021; Kimbrough et al., 2020). This study was underpowered to examine the influence of sex, leading us to combine data from both sexes.

#### Imaging

##### Acquisition

Sample imaging was performed using a Zeiss Lightsheet 7 microscope operated with Zen Black version 3.1 and equipped with a detection objective (Fluar 2.5x/0.12 M27; WD=8.7mm), two illumination objectives (5x/0.1 foc), two PCO.edge 4.2 cameras, and a Mesoscale Imaging System from Translucence (imaging chamber with glass windows, objective adapter, and magnetic sample holders). The posterior portion of each hemisphere was securely mounted to a magnetic sample holder with Loctite Ultra Gel Control Super Glue and submerged into the imaging chamber filled with ethyl cinnamate. Dual side illumination (with online fusion of left and right light sheet images) was used to simultaneously acquire images of autofluorescence (later used for registering samples to the mouse brain atlas as it provides structural information) and IF labeling with separate cameras at 3.53 µm x and y resolution (0.52x zoom), moving the sample 3.5 µm for each z-step to acquire images spanning each hemisphere. Autofluorescence was captured using 488 nm excitation (505-530 nm band pass emission) at 8% of 30 mW laser power, while IF labeling was captured using 638 nm excitation (660 nm long pass emission) at 20% of 75 mW laser power. Exposure time for both excitation wavelengths was 50 ms. Pivot scanning was used to limit image artifacts with light sheet thickness set to 10.61 µm.

##### Pre-processing

Multiple z-stack image tiles (∼2×5) encompassed the hemisphere, with x and y tile dimensions of ∼800 × 688 pixels (limited in size for ∼uniform z resolution) and 10% overlap. An automated stitching function in Zen Blue 3.3 (Zeiss, CH) was used to stitch tiles together according to a mid-z-stack reference image. Computational image analysis was performed on a DELL Precision 7920 Tower, equipped with 40 CPUs, 755 GB of RAM, four 2 TB NVMe SSDs, 24 GB of GPU memory, and Ubuntu 18.04.4. Stitched image volumes (.czi file type) were automatically prepped for registration to the atlas using custom bash scripts (4.4.20(1)) and Fiji (2.1.0/1.53c) (Schindelin et al., 2012) macros. Images were minimally cropped in x and y if there was an uneven number of pixels and saved as a tif series.

To improve registration, the display range of the autofluorescence tif series was linearly adjusted to zero most voxels outside tissue (otherwise external voxels can pull atlas labels outward). The autofluorescence tif series was downsampled by a factor of 2, reoriented (to match the averaged template), and converted to a .nii.gz file type with miracl_conv_convertTIFFtoNII.py from MIRACL (downloaded 05/20/2020; https://miracl.readthedocs.io/en/latest/) (Goubran et al., 2019). When hemisections were imprecise, excess medial tissue was digitally trimmed or missing contralateral tissue was added using 3D Slicer (Fedorov et al., 2012) (4.11 with the SegmentEditorExtraEffects extension installed for “Surface Cut” and “Mask Volume” tools). The intensity of damaged or dim tissue was adjusted in a few cases to fine tune intensity matching between the tissue and template during registration. For intensity-based registration, we modified miracl_reg_clar-allen_whole_brain.sh (Goubran et al., 2019) to work with an LSFM-acquired average template brain (Perens, Salinas, et al., 2021) in alignment with the common coordinate framework version 3 of the Allen brain atlas (ABA) (Wang et al., 2020). We resampled the template and atlas to 25 µm isotropic resolution (bicubic interpolation for the template and nearest neighbor interpolation for atlas labels), removed the olfactory bulb, and lowered the intensity of high intensity atlas region IDs (to better match ABA label coding and constrict the display range for better viewing).

MIRACL’s registration script uses tools from ANTs for down-sampling the autofluorescence image and aligning it with the template. Initial alignment with antsAffineInitializer involves several iterations of search configurations (Goubran et al., 2019). Alignment is refined via intensity-based b-spline transformations (rigid, affine, and then deformable) with 4 levels of resolution (Goubran et al., 2019). Twenty-five µm LSFM ABA labels are then warped to native space (10 µm isotropic resolution) and a corresponding autofluorescence image is output (Goubran et al., 2019). These images enable visual inspection of registration accuracy using ITK-SNAP (Yushkevich et al., 2006) (3.6.0), aided by a custom LUT for the atlas (applying ABA coloring [Figure 1C] and allowing users to hover the cursor to see region names).

To enhance the sensitivity of voxel-wise analysis, autofluorescence was removed from IF image volumes using rolling ball background subtraction in FIJI (Sternberg, 1983) (radius=4 pixels). Background subtracted tifs were prepped for warping to 25 µm resolution atlas space using miracl_conv_convertTIFFtoNII.py, warped using miracl_reg_warp_clar_data_to_gubra.sh as well as the transformation matrices and deformation fields obtained from registration of autofluorescence data (Goubran et al., 2019), and z-scored. These transformations served to normalize both the position of all anatomical regions along common coordinates (registration) and the fluorescence intensity to control for variations in clearing efficiency, immunolabeling, and image acquisition (Carvajal-Camelo et al., 2021).

##### Statistics and brain sample criteria

Sample sizes (number of mice) are summarized in the Animals section and in Table S2. They can also be viewed for specific experiments via individual data points in cluster validation bar graphs. Sample sizes for activity mapping were comparable to other published work (DeNardo et al., 2019; Kimbrough et al., 2020). Statistical tests and comparisons are described in detail in this section but are also mentioned in results and/or figure legends. To control for multiple comparisons with voxel-wise analyses, we used FDR correction because it is an accepted method of controlling for false positives in fMRI data (Benjamini & Hochberg, 1995; Genovese et al., 2002; Perens, Skytte, et al., 2021). For *post hoc* comparisons of c-Fos^+^ cell density, we assumed that on a population level, density measurements would be normally distributed; hence, normality was not verified. Thresholds and criteria for significance for each test are described in this section. Mean ± SEM are shown. Mice were randomly assigned to groups with the goal of having similar sample sizes for c-Fos mapping, with the constraint that available littermates were kept together and exposed to the same conditions. Brain samples were excluded prior to analysis due to tissue damage, poor registration quality, genotype, or blurring of IF labeling from glue used to affix the posterior brain to a sample-holder for imaging, thus preventing accurate cell segmentation. Clusters identified in the hindbrain were excluded from analysis due to artifactual image blurring. Left hemispheres were primarily used for anti-c-Fos immunolabeling, whereas right hemispheres were primarily used for anti-RFP immunolabeling. Two right hemispheres were used for anti-c-Fos immunolabeling, and one left hemisphere was used for anti-RFP immunolabeling.

##### Voxel-wise statistics

Voxel-wise statistics were performed using FSL (Smith et al., 2004) (FMRIB Software Library 6.0.2:a4f562d9; www.fmrib.ox.ac.uk/fsl). After background subtraction, c-Fos-IF image volumes were warped to 25 µm atlas space, z-score normalized using a hemispheric template mask and visually compared with a wire frame version of the atlas to confirm acceptable alignment. Normalized c-Fos-IF images were merged into a 4D file with fslmerge and smoothed with a 50 µm full width at half maximum Gaussian filter. Generalized linear modeling was performed using the randomise_parallel tool in FSL according to a two-way ANOVA design and compared to a null distribution built by nonparametric permutation testing (Winkler et al., 2014) (6,000 permutations). Voxelized F-contrast outputs were subjected to multiple comparisons correction by false discovery rate (FDR) using a hemisphere mask lacking the cerebellum and thresholded to preserve clusters ≥100 voxels (≥1.15625 × 10^-3^ mm^3^), limiting the occurrence of false positive voxels (Benjamini & Hochberg, 1995; Bennett et al., 2009; Genovese et al., 2002). To identify an optimal correction threshold for FDR (q value), multiple q values were used to threshold main effect and interaction maps, followed by determining the rate of true positives (number of clusters with a difference in c-Fos^+^ cell density divided by the total number of clusters). Thresholds (q values) were selected to maximize specificity and stringency, leading us to emphasize results corresponding to q < 0.01 (p < 0.000167131) for main effects and q < 0.15 (p < 0.000334382) for the interaction. A total of 486 brain regions were included in the analysis, excluding regions with image artifacts (i.e., the cerebellum, medulla, and small portions of the midbrain and visual cortex) and the excised olfactory bulb.

##### c-Fos^+^ cell segmentation

Using the pixel classification workflow in Ilastik (1.3.3) (Berg et al., 2019) and three training images from three samples of each condition, pixels belonging to background and either c-Fos^+^ or tdTomato^+^ cells were classified. After initial training with all image features, 7 optimal image features were automatically selected for training the random forest classifier and used during training by five independent raters. Custom scripts were used to run Ilastik in headless mode and generate binary cell segmentations for all samples and raters. In addition to using an automated cell classification algorithm based on sparse-user input to control for subjective cell classification, we also controlled for variability between raters by preserving voxels classified as c-Fos^+^ or tdTomato^+^ cells if identified by ≥ 3 raters, thus generating consensus images of cell segmentation.

##### Cluster validation

To determine effect direction, and if clusters represented valid differences in immunolabeled cell density, each cluster surviving FDR-correction and extent-thresholding was subjected to *post hoc* tests of cell density comparisons (α = 0.05). For this, clusters were warped to tissue space for each sample using nearest neighbor interpolation, upsampled to full resolution, binarized and cropped based on their spatial extent. The binary cluster mask was multiplied by the corresponding region of the 3D consensus segmentation image to zero-out voxels beyond the cluster bounds. Cell counting was accelerated by using the GPU via a CLIJx plugin (0.30.1.16) for Fiji. Cluster volumes were quantified with fslstats using the native resolution binary cluster mask for cell density calculations. GraphPad Prism9 or a custom R script (version 4.1.3) was used to perform for *post hoc* tests, including unpaired t-tests for main effect clusters and 2×2 ANOVAs with Tukey’s post-test comparisons for the interaction clusters.

For volumetric regional analysis within clusters, the warped atlas was scaled to full resolution in tissue space and multiplied by the cluster mask to obtain regional volumes (ABA regions were coded by voxel intensities). Regional volumes for all hierarchal levels were calculated with custom Excel templates and a user-defined function in Visual Basic for Applications was used for color coding tables based on the ABA. Interactive sunburst plots summarizing the regional volumes of clusters in atlas space were generated with a custom python script and plotted using Flourish (2022). Three-dimensional model videos of valid clusters were made with step T3 in DSI-Studio (Aug., 24^th^, 2022), using the 25 µm binarized atlas as a full isosurface, clusters as regions, and a RGBA txt file for region colors. Videos were organized and exported with Adobe Premiere Pro (2022).

For qualitative validation (Figure S1), the thresholded FDR-adjusted statistical map was warped to native space for each sample and cropped to each cluster’s extent. The slice with the greatest integrated density of significant voxels was identified to extract montage tiles for c-Fos-IF, rolling ball subtracted images, and consensus cell segmentations. The cluster perimeter was drawn onto each tile using a Fiji macro. Since montage tile sizes differ due to warping, average tile dimensions were determined to uniformly size tiles. ImageMagick (7.1.0-43) was used to determine dimensions of .tif slices and to make montages. During the automated workflow, image volumes were cropped to the bounding box of each cluster to speed up processing, limit RAM usage and file sizes, as well as to restrict x and y dimensions for montages. Additionally, bit depth was reduced when possible and files were generally saved as compressed .nii.gz files.

##### Co-activity analysis

To examine potential functional relationships among clusters representing validated effects of psilocybin on the c-Fos^+^ cell density, we calculated the inter-mouse Pearson correlation of c-Fos^+^ cell densities between each pair of valid clusters, generating a 27 × 27 correlation matrix for each condition. To note, this procedure of calculating activity correlations between regions resembles fMRI methods to determine functional connectivity. Here we refer to such correlations as “co-activity” to avoid confusion with “connectivity” in the fMRI literature. To examine changes in co-activity, we subtracted the saline correlations from those of psilocybin. Correlation matrices were calculated in R (version 4.1.3) using the psych package (version 2.2.3) and were visualized using the corrplot package (version 0.92).

Hierarchical clustering was used to delineate modules of co-active clusters in each correlation matrix. The correlation coefficient between each cluster pair was converted to the complete Euclidean distance for clustering. The derived dendrograms were trimmed at different tree heights (between 30–100% of the maximal tree height), to compare the total number of modules across hierarchical levels. A reliable difference in modularity should be consistent and robust between groups across different tree-cutting thresholds. We chose to highlight modularity results cut at 60% of the maximum tree height, which is near the elbow of the height by modularity curve. Hierarchical clustering was conducted using the built-in hclust function in the R stats package and was visualized using the heatmap.2 function in gplots (version 3.1.3).

Inspired by work from Kimbrough et al. (Kimbrough et al., 2020, 2021), looking at inter- and intra-module characteristics of between-region co-activity as defined by c-Fos^+^ cell counts, we used the same graph theory approach to examine psilocybin-induced changes in regional centrality metrics, including the Z-scored version of within-module degree (WMDz) and the participation coefficient (PC). The WMDz represents the relative importance of a cluster within its own module, which corresponds to intra-module connectivity. The PC measures the extent a cluster correlates with multiple modules, which corresponds to the inter-module connectivity (Guimerà & Nunes Amaral, 2005). As recommended elsewhere (Kimbrough et al., 2020, 2021) we thresholded the co-activity correlograms by removing co-activity edges weaker than r = 0.75 and excluded negative co-activity edges following conventions as no consensus was reached regarding how to handle them in graph theory analysis (Hallquist & Hillary, 2018). Graph theory analysis was conducted using the Brain Connectivity Toolbox (Rubinov & Sporns, 2010) in Matlab R2020b.

## Data and code availability

Raw data (∼30 GB/hemisphere) is stored locally on external hard drives and on a remote RAID-enabled Synology server in the Forsythe Hall Data Center at Stanford and will be made available by the lead contact upon reasonable request.

Scripts making up UNRAVEL, which we used to automate analysis of LSFM data, are publicly available at: github.com/b-heifets/. This site has our detailed protocols regarding TRAPing, sample prep, IF staining, iDISCO+ clearing, LSFM imaging, and data analysis. Please contact Boris Heifets if you have questions and/or suggestions.

## References

Ali, A. E. A., Wilson, Y. M., & Murphy, M. (2009). A single exposure to an enriched environment stimulates the activation of discrete neuronal populations in the brain of the fos-tau-lacZ mouse. Neurobiology of Learning and Memory, 92(3), 381–390. https://doi.org/10.1016/j.nlm.2009.05.004

Allen, W. E., DeNardo, L. A., Chen, M. Z., Liu, C. D., Loh, K. M., Fenno, L. E., Ramakrishnan, C., Deisseroth, K., & Luo, L. (2017). Thirst-associated preoptic neurons encode an aversive motivational drive. Science, 357(6356), 1149–1155. https://doi.org/10.1126/science.aan6747

Baroncelli, L., Braschi, C., Spolidoro, M., Begenisic, T., Sale, A., & Maffei, L. (2010). Nurturing brain plasticity: Impact of environmental enrichment. Cell Death & Differentiation, 17(7), Article 7. https://doi.org/10.1038/cdd.2009.193

Benjamini, Y., & Hochberg, Y. (1995). Controlling the False Discovery Rate: A Practical and Powerful Approach to Multiple Testing. Journal of the Royal Statistical Society: Series B (Methodological), 57(1), 289–300. https://doi.org/10.1111/j.2517-6161.1995.tb02031.x

Bennett, C., Miller, M., & Wolford, G. (2009). Neural correlates of interspecies perspective taking in the post-mortem Atlantic Salmon: An argument for multiple comparisons correction. NeuroImage, 47, S125. https://doi.org/10.1016/S1053-8119(09)71202-9

Berg, S., Kutra, D., Kroeger, T., Straehle, C. N., Kausler, B. X., Haubold, C., Schiegg, M., Ales, J., Beier, T., Rudy, M., Eren, K., Cervantes, J. I., Xu, B., Beuttenmueller, F., Wolny, A., Zhang, C., Koethe, U., Hamprecht, F. A., & Kreshuk, A. (2019). ilastik: Interactive machine learning for (bio)image analysis. Nature Methods, 16(12), 1226–1232. https://doi.org/10.1038/s41592-019-0582-9

Carhart-Harris, R. L. (2023). Translational Challenges in Psychedelic Medicine. New England Journal of Medicine, 388(5), 476–477. https://doi.org/10.1056/NEJMcibr2213109

Carhart-Harris, R. L., Bolstridge, M., Day, C. M. J., Rucker, J., Watts, R., Erritzoe, D. E., Kaelen, M., Giribaldi, B., Bloomfield, M., Pilling, S., Rickard, J. A., Forbes, B., Feilding, A., Taylor, D., Curran, H. V., & Nutt, D. J. (2018). Psilocybin with psychological support for treatment-resistant depression: Six-month follow-up. Psychopharmacology, 235(2), 399–408. https://doi.org/10.1007/s00213-017-4771-x

Carhart-Harris, R. L., Erritzoe, D., Williams, T., Stone, J. M., Reed, L. J., Colasanti, A., Tyacke, R. J., Leech, R., Malizia, A. L., Murphy, K., Hobden, P., Evans, J., Feilding, A., Wise, R. G., & Nutt, D. J. (2012). Neural correlates of the psychedelic state as determined by fMRI studies with psilocybin. Proceedings of the National Academy of Sciences, 109(6), 2138–2143. https://doi.org/10.1073/pnas.1119598109

Carvajal-Camelo, E. E., Bernal, J., Oliver, A., Lladó, X., Trujillo, M., & Initiative, T. A. D. N. (2021). Evaluating the Effect of Intensity Standardisation on Longitudinal Whole Brain Atrophy Quantification in Brain Magnetic Resonance Imaging. Applied Sciences, 11(4), Article 4. https://doi.org/10.3390/app11041773

Chowdhury, A., & Caroni, P. (2018). Time units for learning involving maintenance of system-wide cFos expression in neuronal assemblies. Nature Communications, 9(1), Article 1. https://doi.org/10.1038/s41467-018-06516-3

Cullen, P. K., Gilman, T. L., Winiecki, P., Riccio, D. C., & Jasnow, A. M. (2015). Activity of the anterior cingulate cortex and ventral hippocampus underlie increases in contextual fear generalization. Neurobiology of Learning and Memory, 124, 19–27. https://doi.org/10.1016/j.nlm.2015.07.001

Davis, A. K., Barrett, F. S., May, D. G., Cosimano, M. P., Sepeda, N. D., Johnson, M. W., Finan, P. H., & Griffiths, R. R. (2021). Effects of Psilocybin-Assisted Therapy on Major Depressive Disorder: A Randomized Clinical Trial. JAMA Psychiatry, 78(5), 481–489. https://doi.org/10.1001/jamapsychiatry.2020.3285

Davoudian, P. A., Shao, L.-X., & Kwan, A. C. (2023). Shared and Distinct Brain Regions Targeted for Immediate Early Gene Expression by Ketamine and Psilocybin. ACS Chemical Neuroscience, acschemneuro.2c00637. https://doi.org/10.1021/acschemneuro.2c00637

DeNardo, L. A., Liu, C. D., Allen, W. E., Adams, E. L., Friedmann, D., Fu, L., Guenthner, C. J., Tessier-Lavigne, M., & Luo, L. (2019). Temporal evolution of cortical ensembles promoting remote memory retrieval. Nature Neuroscience, 22(3), 460–469. https://doi.org/10.1038/s41593-018-0318-7

Eklund, A., Nichols, T. E., & Knutsson, H. (2016). Cluster failure: Why fMRI inferences for spatial extent have inflated false-positive rates. Proceedings of the National Academy of Sciences, 113(28), 7900–7905. https://doi.org/10.1073/pnas.1602413113

Ey, E., Leblond, C. S., & Bourgeron, T. (2011). Behavioral profiles of mouse models for autism spectrum disorders. Autism Research, 4(1), 5–16. https://doi.org/10.1002/aur.175

Fadahunsi, N., Lund, J., Breum, A. W., Mathiesen, C. V., Larsen, I. B., Knudsen, G. M., Klein, A. B., & Clemmensen, C. (2022). Acute and long-term effects of psilocybin on energy balance and feeding behavior in mice. Translational Psychiatry, 12(1), Article 1. https://doi.org/10.1038/s41398-022-02103-9

Fedorov, A., Beichel, R., Kalpathy-Cramer, J., Finet, J., Fillion-Robin, J.-C., Pujol, S., Bauer, C., Jennings, D., Fennessy, F., Sonka, M., Buatti, J., Aylward, S., Miller, J. V., Pieper, S., & Kikinis, R. (2012). 3D Slicer as an image computing platform for the Quantitative Imaging Network. Magnetic Resonance Imaging, 30(9), 1323–1341. https://doi.org/10.1016/j.mri.2012.05.001

Forstmann, M., Yudkin, D. A., Prosser, A. M. B., Heller, S. M., & Crockett, M. J. (2020). Transformative experience and social connectedness mediate the mood-enhancing effects of psychedelic use in naturalistic settings. Proceedings of the National Academy of Sciences of the United States of America, 117(5), 2338–2346. https://doi.org/10.1073/pnas.1918477117

Genovese, C. R., Lazar, N. A., & Nichols, T. (2002). Thresholding of Statistical Maps in Functional Neuroimaging Using the False Discovery Rate. NeuroImage, 15(4), 870–878. https://doi.org/10.1006/nimg.2001.1037

Golden, C. T., & Chadderton, P. (2022). Psilocybin reduces low frequency oscillatory power and neuronal phase-locking in the anterior cingulate cortex of awake rodents. Scientific Reports, 12(1), Article 1. https://doi.org/10.1038/s41598-022-16325-w

Golden, T. L., Magsamen, S., Sandu, C. C., Lin, S., Roebuck, G. M., Shi, K. M., & Barrett, F. S. (2022). Effects of Setting on Psychedelic Experiences, Therapies, and Outcomes: A Rapid Scoping Review of the Literature. Current Topics in Behavioral Neurosciences, 56, 35–70. https://doi.org/10.1007/7854_2021_298

Goodwin, G. M., Aaronson, S. T., Alvarez, O., Arden, P. C., Baker, A., Bennett, J. C., Bird, C., Blom, R. E., Brennan, C., Brusch, D., Burke, L., Campbell-Coker, K., Carhart-Harris, R., Cattell, J., Daniel, A., DeBattista, C., Dunlop, B. W., Eisen, K., Feifel, D., … Malievskaia, E. (2022). Single-Dose Psilocybin for a Treatment-Resistant Episode of Major Depression. New England Journal of Medicine, 387(18), 1637–1648. https://doi.org/10.1056/NEJMoa2206443

Goubran, M., Leuze, C., Hsueh, B., Aswendt, M., Ye, L., Tian, Q., Cheng, M. Y., Crow, A., Steinberg, G. K., McNab, J. A., Deisseroth, K., & Zeineh, M. (2019). Multimodal image registration and connectivity analysis for integration of connectomic data from microscopy to MRI. Nature Communications, 10(1), 5504. https://doi.org/10.1038/s41467-019-13374-0

Gouzoulis-Mayfrank, E., Schreckenberger, M., Sabri, O., Arning, C., Thelen, B., Spitzer, M., Kovar, K.-A., Hermle, L., Büll, U., & Sass, H. (1999). Neurometabolic Effects of Psilocybin, 3,4-Methylenedioxyethylamphetamine (MDE) and d-Methamphetamine in Healthy Volunteers: A Double-Blind, Placebo-Controlled PET Study with [18F]FDG. Neuropsychopharmacology, 20(6), 565–581. https://doi.org/10.1016/S0893-133X(98)00089-X

Guimerà, R., & Nunes Amaral, L. A. (2005). Functional cartography of complex metabolic networks. Nature, 433(7028), Article 7028. https://doi.org/10.1038/nature03288

Hallquist, M. N., & Hillary, F. G. (2018). Graph theory approaches to functional network organization in brain disorders: A critique for a brave new small-world. Network Neuroscience, 3(1), 1–26. https://doi.org/10.1162/netn_a_00054

Hansen, H. H., Perens, J., Roostalu, U., Skytte, J. L., Salinas, C. G., Barkholt, P., Thorbek, D. D., Rigbolt, K. T. G., Vrang, N., Jelsing, J., & Hecksher-Sørensen, J. (2021). Whole-brain activation signatures of weight-lowering drugs. Molecular Metabolism, 47, 101171. https://doi.org/10.1016/j.molmet.2021.101171

Hartogsohn, I. (2017). Constructing drug effects: A history of set and setting. Drug Science, Policy and Law, 3. https://doi.org/10.1177/2050324516683325

Hesselgrave, N., Troppoli, T. A., Wulff, A. B., Cole, A. B., & Thompson, S. M. (2021). Harnessing psilocybin: Antidepressant-like behavioral and synaptic actions of psilocybin are independent of 5-HT2R activation in mice. Proceedings of the National Academy of Sciences of the United States of America, 118(17), e2022489118. https://doi.org/10.1073/pnas.2022489118

Hibicke, M., Landry, A. N., Kramer, H. M., Talman, Z. K., & Nichols, C. D. (2020). Psychedelics, but Not Ketamine, Produce Persistent Antidepressant-like Effects in a Rodent Experimental System for the Study of Depression. ACS Chemical Neuroscience, 11(6), 864–871. https://doi.org/10.1021/acschemneuro.9b00493

Hoffman, G. E., Smith, M. S., & Verbalis, J. G. (1993). C-Fos and Related Immediate Early Gene Products as Markers of Activity in Neuroendocrine Systems. Frontiers in Neuroendocrinology, 14(3), 173–213. https://doi.org/10.1006/frne.1993.1006

Jin, M., Nguyen, J. D., Weber, S. J., Mejias-Aponte, C. A., Madangopal, R., & Golden, S. A. (2022). SMART: An Open-Source Extension of WholeBrain for Intact Mouse Brain Registration and Segmentation. Eneuro, 9(3), ENEURO.0482-21.2022. https://doi.org/10.1523/ENEURO.0482-21.2022

Johnson, M. W., Hendricks, P. S., Barrett, F. S., & Griffiths, R. R. (2019). Classic psychedelics: An integrative review of epidemiology, therapeutics, mystical experience, and brain network function. Pharmacology & Therapeutics, 197, 83–102. https://doi.org/10.1016/j.pharmthera.2018.11.010

Kimbrough, A., Kallupi, M., Smith, L. C., Simpson, S., Collazo, A., & George, O. (2021). Characterization of the Brain Functional Architecture of Psychostimulant Withdrawal Using Single-Cell Whole-Brain Imaging. ENeuro, 8(6). https://doi.org/10.1523/ENEURO.0208-19.2021

Kimbrough, A., Lurie, D. J., Collazo, A., Kreifeldt, M., Sidhu, H., Macedo, G. C., D’Esposito, M., Contet, C., & George, O. (2020). Brain-wide functional architecture remodeling by alcohol dependence and abstinence. Proceedings of the National Academy of Sciences, 117(4), 2149–2159. https://doi.org/10.1073/pnas.1909915117

Kwan, A. C., Olson, D. E., Preller, K. H., & Roth, B. L. (2022). The neural basis of psychedelic action. Nature Neuroscience, 25(11), 1407–1419. https://doi.org/10.1038/s41593-022-01177-4

Lewis, C. R., Preller, K. H., Kraehenmann, R., Michels, L., Staempfli, P., & Vollenweider, F. X. (2017). Two dose investigation of the 5-HT-agonist psilocybin on relative and global cerebral blood flow. NeuroImage, 159, 70–78. https://doi.org/10.1016/j.neuroimage.2017.07.020

Morgan, J. I., Cohen, D. R., Hempstead, J. L., & Curran, T. (1987). Mapping patterns of c-fos expression in the central nervous system after seizure. Science (New York, N.Y.), 237(4811), 192–197. https://doi.org/10.1126/science.3037702

Nithianantharajah, J., & Hannan, A. J. (2006). Enriched environments, experience-dependent plasticity and disorders of the nervous system. Nature Reviews Neuroscience, 7(9), Article 9. https://doi.org/10.1038/nrn1970

Noble, S., Scheinost, D., & Constable, R. T. (2020). Cluster failure or power failure? Evaluating sensitivity in cluster-level inference. NeuroImage, 209, 116468. https://doi.org/10.1016/j.neuroimage.2019.116468

Nygart, V. A., Pommerencke, L. M., Haijen, E., Kettner, H., Kaelen, M., Mortensen, E. L., Nutt, D. J., Carhart-Harris, R. L., & Erritzoe, D. (2022). Antidepressant effects of a psychedelic experience in a large prospective naturalistic sample. Journal of Psychopharmacology (Oxford, England), 36(8), 932–942. https://doi.org/10.1177/02698811221101061

Perens, J., Salinas, C. G., Skytte, J. L., Roostalu, U., Dahl, A. B., Dyrby, T. B., Wichern, F., Barkholt, P., Vrang, N., Jelsing, J., & Hecksher-Sørensen, J. (2021). An Optimized Mouse Brain Atlas for Automated Mapping and Quantification of Neuronal Activity Using iDISCO+ and Light Sheet Fluorescence Microscopy. Neuroinformatics, 19(3), 433–446. https://doi.org/10.1007/s12021-020-09490-8

Perens, J., Skytte, J. L., Salinas, C. G., Hecksher-Sorensen, J., Dyrby, T. B., & Dahl, A. B. (2021). Comparative Study Of Voxel-Based Statistical Analysis Methods For Fluorescently Labelled And Light Sheet Imaged Whole-Brain Samples. 2021 IEEE 18th International Symposium on Biomedical Imaging (ISBI), 1433–1437. https://doi.org/10.1109/ISBI48211.2021.9434015

Renier, N., Adams, E. L., Kirst, C., Wu, Z., Azevedo, R., Kohl, J., Autry, A. E., Kadiri, L., Umadevi Venkataraju, K., Zhou, Y., Wang, V. X., Tang, C. Y., Olsen, O., Dulac, C., Osten, P., & Tessier-Lavigne, M. (2016). Mapping of Brain Activity by Automated Volume Analysis of Immediate Early Genes. Cell, 165(7), 1789–1802. https://doi.org/10.1016/j.cell.2016.05.007

Renier, N., Wu, Z., Simon, D. J., Yang, J., Ariel, P., & Tessier-Lavigne, M. (2014). iDISCO: A Simple, Rapid Method to Immunolabel Large Tissue Samples for Volume Imaging. Cell, 159(4), 896–910. https://doi.org/10.1016/j.cell.2014.10.010

Roseman, L., Nutt, D. J., & Carhart-Harris, R. L. (2018). Quality of Acute Psychedelic Experience Predicts Therapeutic Efficacy of Psilocybin for Treatment-Resistant Depression. Frontiers in Pharmacology, 8. https://www.frontiersin.org/articles/10.3389/fphar.2017.00974

Rubinov, M., & Sporns, O. (2010). Complex network measures of brain connectivity: Uses and interpretations. NeuroImage, 52(3), 1059–1069. https://doi.org/10.1016/j.neuroimage.2009.10.003

Sagar, S. M., Sharp, F. R., & Curran, T. (1988). Expression of c-fos protein in brain: Metabolic mapping at the cellular level. Science (New York, N.Y.), 240(4857), 1328–1331. https://doi.org/10.1126/science.3131879

Schindelin, J., Arganda-Carreras, I., Frise, E., Kaynig, V., Longair, M., Pietzsch, T., Preibisch, S., Rueden, C., Saalfeld, S., Schmid, B., Tinevez, J.-Y., White, D. J., Hartenstein, V., Eliceiri, K., Tomancak, P., & Cardona, A. (2012). Fiji: An open-source platform for biological-image analysis. Nature Methods, 9(7), 676–682. https://doi.org/10.1038/nmeth.2019

Schmack, K., Bosc, M., Ott, T., Sturgill, J. F., & Kepecs, A. (2021). Striatal dopamine mediates hallucination-like perception in mice. Science, 372(6537), eabf4740. https://doi.org/10.1126/science.abf4740

Sert, N. P. du, Hurst, V., Ahluwalia, A., Alam, S., Avey, M. T., Baker, M., Browne, W. J., Clark, A., Cuthill, I. C., Dirnagl, U., Emerson, M., Garner, P., Holgate, S. T., Howells, D. W., Karp, N. A., Lazic, S. E., Lidster, K., MacCallum, C. J., Macleod, M., … Würbel, H. (2020). The ARRIVE guidelines 2.0: Updated guidelines for reporting animal research. PLOS Biology, 18(7), e3000410. https://doi.org/10.1371/journal.pbio.3000410

Sgambato, V., Abo, V., Rogard, M., Besson, M. J., & Deniau, J. M. (1997). Effect of electrical stimulation of the cerebral cortex on the expression of the fos protein in the basal ganglia. Neuroscience, 81(1), 93–112. https://doi.org/10.1016/S0306-4522(97)00179-6

Shao, L.-X., Liao, C., Gregg, I., Davoudian, P. A., Savalia, N. K., Delagarza, K., & Kwan, A. C. (2021). Psilocybin induces rapid and persistent growth of dendritic spines in frontal cortex in vivo. Neuron, 109(16), 2535-2544.e4. https://doi.org/10.1016/j.neuron.2021.06.008

Smith, S. M., Jenkinson, M., Woolrich, M. W., Beckmann, C. F., Behrens, T. E. J., Johansen-Berg, H., Bannister, P. R., De Luca, M., Drobnjak, I., Flitney, D. E., Niazy, R. K., Saunders, J., Vickers, J., Zhang, Y., De Stefano, N., Brady, J. M., & Matthews, P. M. (2004). Advances in functional and structural MR image analysis and implementation as FSL. NeuroImage, 23, S208–S219. https://doi.org/10.1016/j.neuroimage.2004.07.051

Solinas, M., Chauvet, C., Thiriet, N., El Rawas, R., & Jaber, M. (2008). Reversal of cocaine addiction by environmental enrichment. Proceedings of the National Academy of Sciences, 105(44), 17145–17150. https://doi.org/10.1073/pnas.0806889105

Sternberg. (1983). Biomedical Image Processing. Computer, 16(1), 22–34. https://doi.org/10.1109/MC.1983.1654163

Torregrossa, M. M., Gordon, J., & Taylor, J. R. (2013). Double Dissociation between the Anterior Cingulate Cortex and Nucleus Accumbens Core in Encoding the Context versus the Content of Pavlovian Cocaine Cue Extinction. Journal of Neuroscience, 33(19), 8370–8377. https://doi.org/10.1523/JNEUROSCI.0489-13.2013

van Praag, H., Kempermann, G., & Gage, F. H. (2000). Neural consequences of enviromental enrichment. Nature Reviews Neuroscience, 1(3), Article 3. https://doi.org/10.1038/35044558

Voelkl, B., Altman, N. S., Forsman, A., Forstmeier, W., Gurevitch, J., Jaric, I., Karp, N. A., Kas, M. J., Schielzeth, H., Van de Casteele, T., & Würbel, H. (2020). Reproducibility of animal research in light of biological variation. Nature Reviews Neuroscience, 21(7), Article 7. https://doi.org/10.1038/s41583-020-0313-3

Vollenweider, F. X. (1998). Advances and Pathophysiological Models of Hallucinogenic Drug Actions in Humans: A Preamble to Schizophrenia Research. Pharmacopsychiatry, 31(S 2), 92–103. https://doi.org/10.1055/s-2007-979353

Vollenweider, F. X., Leenders, K. L., Scharfetter, C., Maguire, P., Stadelmann, O., & Angst, J. (1997). Positron Emission Tomography and Fluorodeoxyglucose Studies of Metabolic Hyperfrontality and Psychopathology in the Psilocybin Model of Psychosis. Neuropsychopharmacology, 16(5), Article 5. https://doi.org/10.1016/S0893-133X(96)00246-1

Wang, Q., Ding, S.-L., Li, Y., Royall, J., Feng, D., Lesnar, P., Graddis, N., Naeemi, M., Facer, B., Ho, A., Dolbeare, T., Blanchard, B., Dee, N., Wakeman, W., Hirokawa, K. E., Szafer, A., Sunkin, S. M., Oh, S. W., Bernard, A., … Ng, L. (2020). The Allen Mouse Brain Common Coordinate Framework: A 3D Reference Atlas. Cell, 181(4), 936-953.e20. https://doi.org/10.1016/j.cell.2020.04.007

Williams, L. M. (2017). Defining biotypes for depression and anxiety based on large-scale circuit dysfunction: A theoretical review of the evidence and future directions for clinical translation. Depression and Anxiety, 34(1), 9–24. https://doi.org/10.1002/da.22556

Wilson, D. I. G., Langston, R. F., Schlesiger, M. I., Wagner, M., Watanabe, S., & Ainge, J. A. (2013). Lateral entorhinal cortex is critical for novel object-context recognition. Hippocampus, 23(5), 352–366. https://doi.org/10.1002/hipo.22095

Winkler, A. M., Ridgway, G. R., Webster, M. A., Smith, S. M., & Nichols, T. E. (2014). Permutation inference for the general linear model. NeuroImage, 92, 381–397. https://doi.org/10.1016/j.neuroimage.2014.01.060

Woo, C.-W., Krishnan, A., & Wager, T. D. (2014). Cluster-extent based thresholding in fMRI analyses: Pitfalls and recommendations. NeuroImage, 91, 412–419. https://doi.org/10.1016/j.neuroimage.2013.12.058

Yushkevich, P. A., Piven, J., Hazlett, H. C., Smith, R. G., Ho, S., Gee, J. C., & Gerig, G. (2006). User-guided 3D active contour segmentation of anatomical structures: Significantly improved efficiency and reliability. NeuroImage, 31(3), 1116–1128. https://doi.org/10.1016/j.neuroimage.2006.01.015

