## Supplemental information for "UNRAVELing the synergistic effects of psilocybin and environment on brain-wide immediate early gene expression in mice"

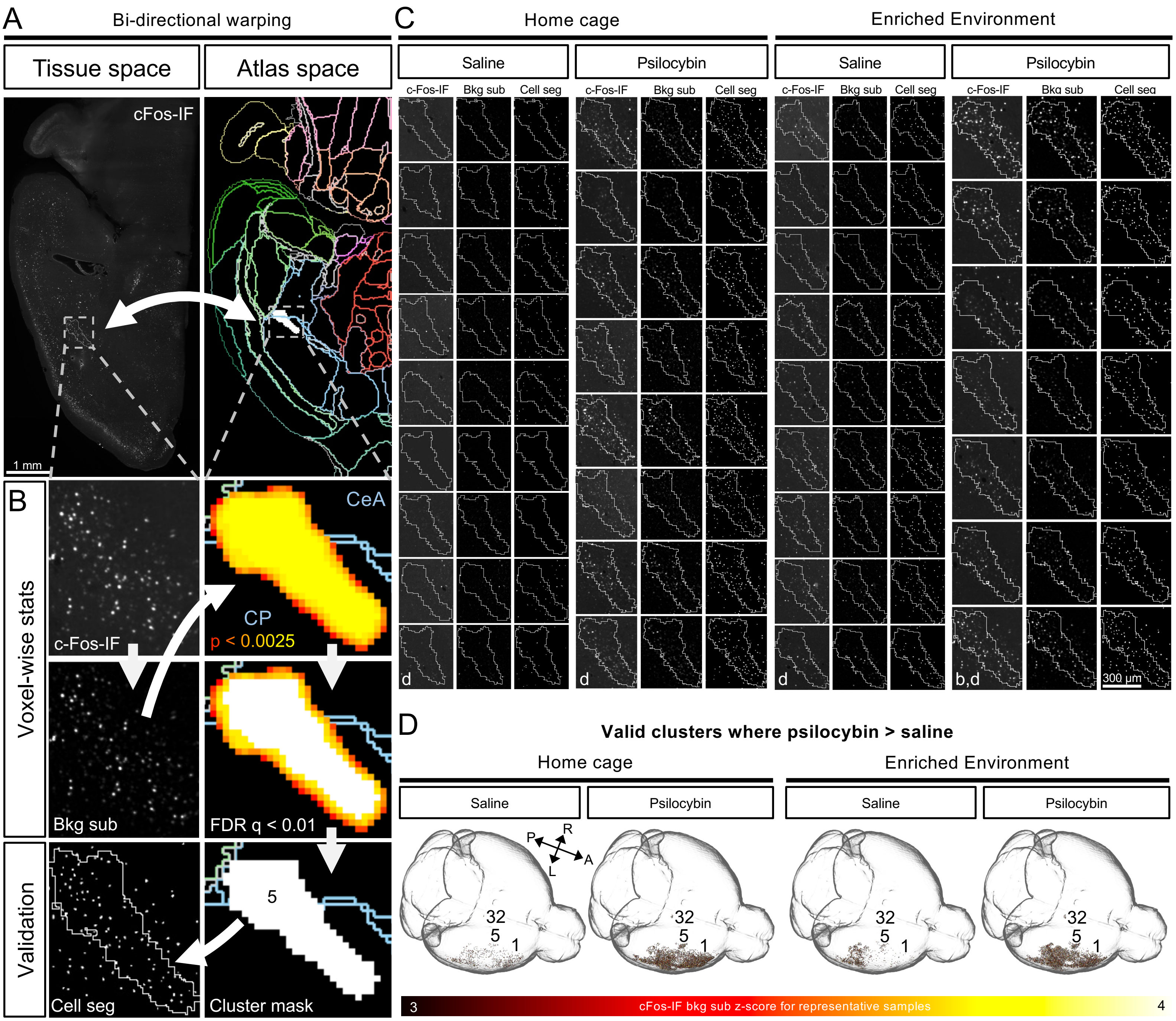

A
Bi-directional warping
Tissue space
Atlas space
cFos-IF
1 mm
B
Voxel-wise stats
c-Fos-IF
Bkg sub
CeA
CP
p < 0.0025
FDR q < 0.01
Validation
Cell seg
5
Cluster mask
C
Home cage
Enriched Environment
Saline
Psilocybin
c-Fos-IF
Bkg sub
Cell seg
d
b,d
300 µm
D
Valid clusters where psilocybin > saline
Home cage
Enriched Environment
Saline
Psilocybin
P
R
L
A
32
5
1
3
cFos-IF bkg sub z-score for representative samples
4

**Figure S1: Example data transformations and qualitative validation.** A) Registration of autofluorescence image volumes to the average template brain and corresponding LSFM version of the Allen brain atlas (ABA) enabled bidirectional warping between tissue space and atlas space. B) c-Fos-IF was background subtracted and warped to LSFM atlas space for voxel-wise analysis. Uncorrected statistical maps were thresholded by FDR correction and cluster extent to define clusters (white) which were warped to tissue space for quantitative validation via cluster-specific cell counting and volume measurements. CeA = central amygdala. CP = caudoputamen. C) Qualitative validation showing montages of raw c-Fos-IF, background subtracted images (Bkg sub), and consensus cell segmentations (Cell seg). Images represent the most significant slice for each cluster and condition. The cluster perimeter is shown for each tile. The row for tile “b” is shown in B. Rows labeled with “d” indicate representative samples used for D. D) MRIcroGL (1.2.20220720b) 3D visualization of background subtracted and z-scored ($z > 3$) cFos-IF in clusters with valid psilocybin-mediated activity increases. The transparent brain is a binary mask of the 25 µm atlas. Select clusters are numerically labeled.

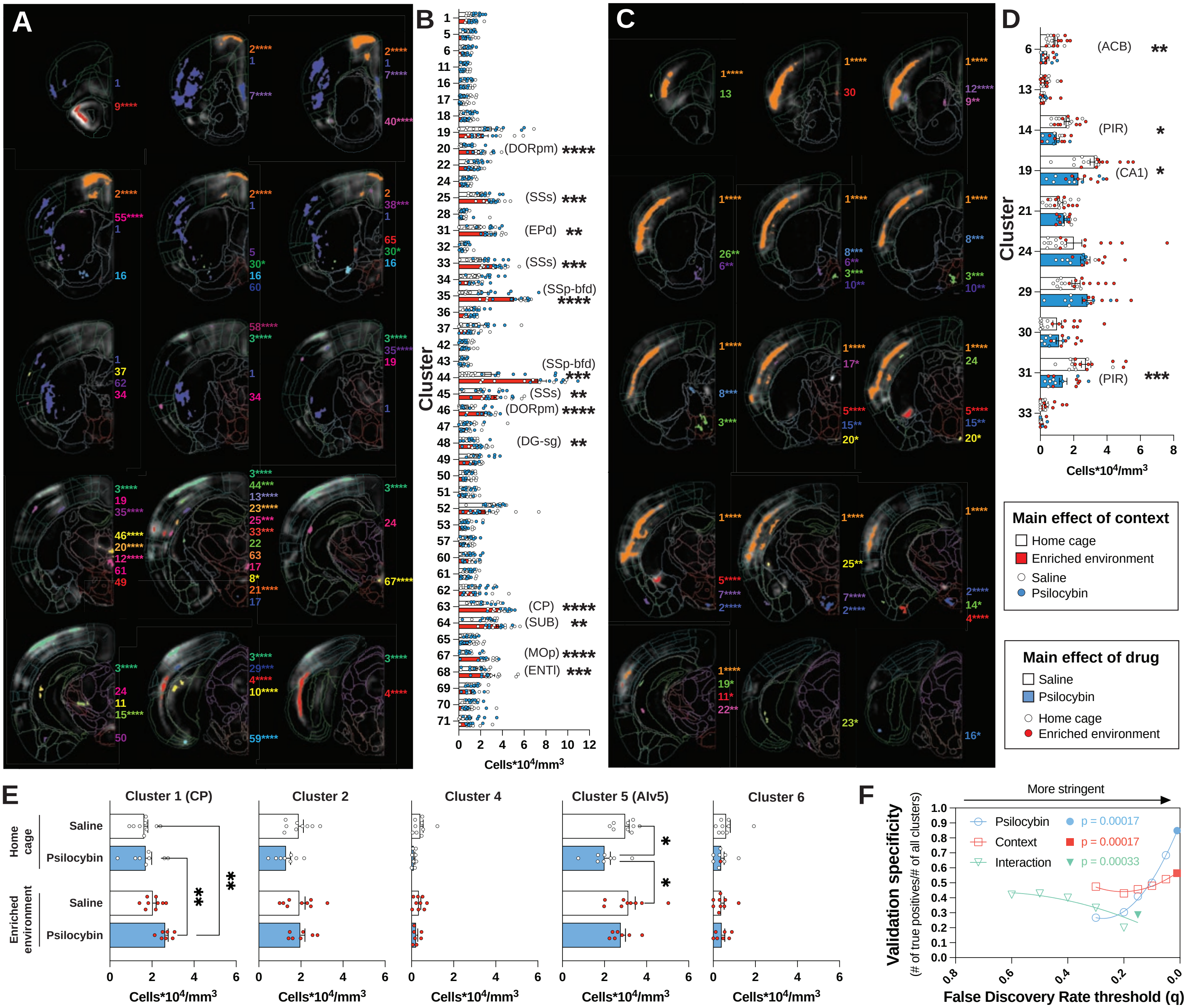
A
B
Cluster
Cells*10^4/mm^3
(DORpm) ****
(SSs) ***
(EPd) **
(SSs) ***
(SSp-bfd) ****
(SSp-bfd) ***
(SSs) **
(DORpm) ****
(DG-sg) **
(CP) ****
(SUB) **
(MOp) ****
(ENTl) ***
C
D
(ACB) **
(PIR) *
(CA1) *
(PIR) ***
Main effect of context
Home cage
Enriched environment
Saline
Psilocybin
Main effect of drug
Saline
Psilocybin
Home cage
Enriched environment
E
Cluster 1 (CP)
Cluster 2
Cluster 4
Cluster 5 (Alv5)
Cluster 6
Home cage
Enriched environment
Saline
Psilocybin
F
More stringent
Validation specificity
(# of true positives/# of all clusters)
False Discovery Rate threshold (q)
Psilocybin p = 0.00017
Context p = 0.00017
Interaction p = 0.00033

**Figure S2: Validation of additional clusters surviving voxel-wise FDR correction (continuation of Figures 2 and 3).** Coronal hemisphere slices illustrating the distribution of clusters surviving FDR correction ($q < 0.01$) from the main effect of C) context and C) psilocybin-treatment. Clusters were randomly colored, and the numerical cluster identity (largest to smallest) is shown along the midline. Clusters overlay a LSFM atlas color coded to Allen atlas labels atop a smoothed, z-score difference map (EE–home cage; psilocybin–saline) depicting the average difference in c-Fos-IF intensity (greyscale). Asterisks report significance from *post hoc* analyses shown in B and D as well as in Figure 2C. *Post hoc* unpaired t-tests of c-Fos$^{+}$ cell density in clusters from the main effect of B) context and D) psilocybin. The primary component region of each valida cluster is shown in parenthesis. E) No *post hoc* 2x2 ANOVA interactions were observed for interaction clusters shown here. Tukey's *post hoc* comparisons of c-Fos$^{+}$ cell density measurements within each cluster indicate validated differences. F) Validation curve for statistical contrasts obtained from the voxel-wise 2x2 ANOVA. Results were FDR corrected across a range of cluster-defining correction thresholds (q) and the corresponding rate of true positives for each set of clusters was determined via cell density measurements and *post hoc* unpaired t-tests. Solid data points, corresponding to the most stringent adjusted p value cut off, represent data presented in this paper. ACB = nucleus accumbens; CA1 = field CA1 ; DG-sg = dentate gyrus, granule cell layer; DORpm = thalamus, polymodal association cortex related; ENTl = entorhinal area, lateral part; EPd = endopiriform nucleus, dorsal part; MOp = primary motor area; p-bfd = primary barrel field; PIR = piriform area; PVR = periventricular region; s = supplemental; SCH = suprachiasmatic nucleus; SS = somatosensory area; SUB = subiculum. *$p < 0.05$, ** $p < 0.01$, ***$p < 0.001$, ****$p < 0.001$.

**Table S1: Valid cluster volumes, locations, and regional composition.** Asterisks indicate significance from *post hoc* unpaired t-tests for main effect clusters in Figure 2 and Figure S2 as well as significance of interactions from *post hoc* 2x2 ANOVAs in Figure 3. For each cluster, grouped by anatomical hierarchy and the effect direction determined by *post hoc* tests, the top 4 subregions are shown (subregions collapsed into parent regions as needed). Total cluster volume is presented as $mm^3$. The center of gravity (CoG) is presented as x,y,z coordinates in the LSFM version of the Allen brain atlas (ABA) used in this paper. Percentages represent the proportion of total cluster volume a brain region occupies. Cells are color coded to region-specific RGB values from the ABA. Ventricles, white matter, the olfactory bulb, medulla, and cerebellum were excluded. a = anterior; bfd = barrel field; d = dorsal; mo = molecular layer; p = primary; l = lateral; ll = lower limb; s = secondary; v = ventral; sg = granular cell layer; po = polymorph layer; white text = other. *$p < 0.05$, ** $p < 0.01$, ***$p < 0.001$, ****$p < 0.001$.

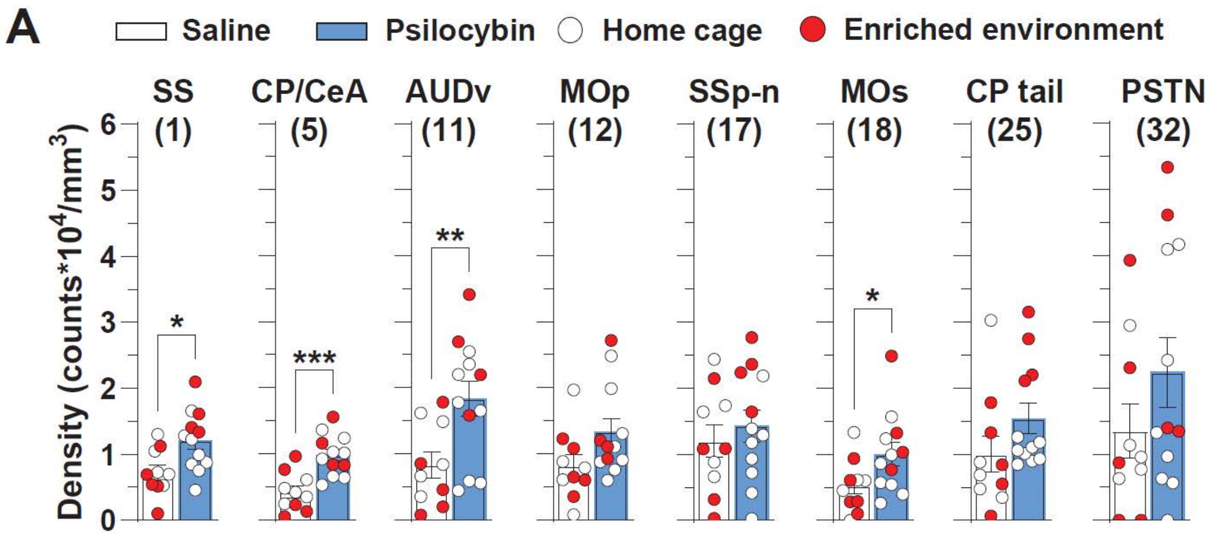


**Figure S3**: **Confirmation of genetic access to ensembles activated by psilocybin (related to Figure 1).** Cluster numbers and the primary component brain region are shown above each graph. $*p < 0.05$, $** \ p < 0.01$, $***p < 0.001$.

**Table S2: Cluster validation summary (related to Figures 2, 3, and S2).** This Excel file summarizes cell densities for all clusters and contrasts, sample sizes, p values from *post hoc* comparisons.

**Video S1. 3D view of valid clusters from main effects ($q < 0.01$) and interactions ($q < 0.15$) colored based on the ABA (related to Figures 2, 3, and S2).**
